# Accounting for DNA Recovery and Cell Culturability Enhances Quantitative Compatibility of Molecular and Legiolert Assays for *Legionella pneumophila*

**DOI:** 10.64898/2026.07.19.739452

**Authors:** Jinhao Yang, Huanqi He, Samantha DiLoreto, Ashwin S Sudarshan, Katherine E. Graham, Linden Neal, Joe Brown, Kelsey J. Pieper, Aron Stubbins, Christopher A. Impellitteri, Ching-Hua Huang, Ameet Pinto

**Author notes:** Corresponding Author: Ameet J. Pinto.

## Abstract

Disagreement between molecular and culture-based assays for *Legionella pneumophila* detection is widely reported, yet comparisons have largely been based on direct assay-derived concentrations or binary positive/negative outcomes. However, it remains unclear whether molecular-culture disagreement reflects concentration-level incompatibility or unaccounted methodological and physiological differences related to DNA recovery and cell culturability. In this study, we also observed disagreement between molecular and Legiolert assays in source and finished drinking water samples collected from eight full-scale drinking water systems across the United States. Molecular thresholds adjusted for DNA recovery and cell culturability only partially resolved these discrepancies. We therefore developed a probabilistic Monte Carlo framework that incorporates sample-specific DNA recovery and cell culturability to evaluate the quantitative consistency of culturable *L. pneumophila* concentrations estimated by molecular and Legiolert assays. Quantitatively consistent and inconsistent samples occurred across both binary concordant and discordant classifications, demonstrating that positive/negative agreement poorly reflects concentration-level comparability. Overall, molecular and Legiolert assays showed strong quantitative consistency once sample-specific DNA recovery and cell culturability were considered. A small proportion of persistent inconsistencies at specific sampling sites, coupled with atypical microbial indicators, suggest that sample heterogeneity likely contributed to the remaining discrepancies. These findings demonstrate that integrating DNA recovery and cell culturability enhanced quantitative consistency between molecular and Legiolert assays and supports the use of molecular methods as rapid quantitative tools to complement culture-based *L. pneumophila* monitoring.

**Synopsis:** Accounting for DNA recovery and cell culturability revealed broad quantitative compatibility between molecular and Legiolert assays for *Legionella pneumophila* in drinking water.

TOC

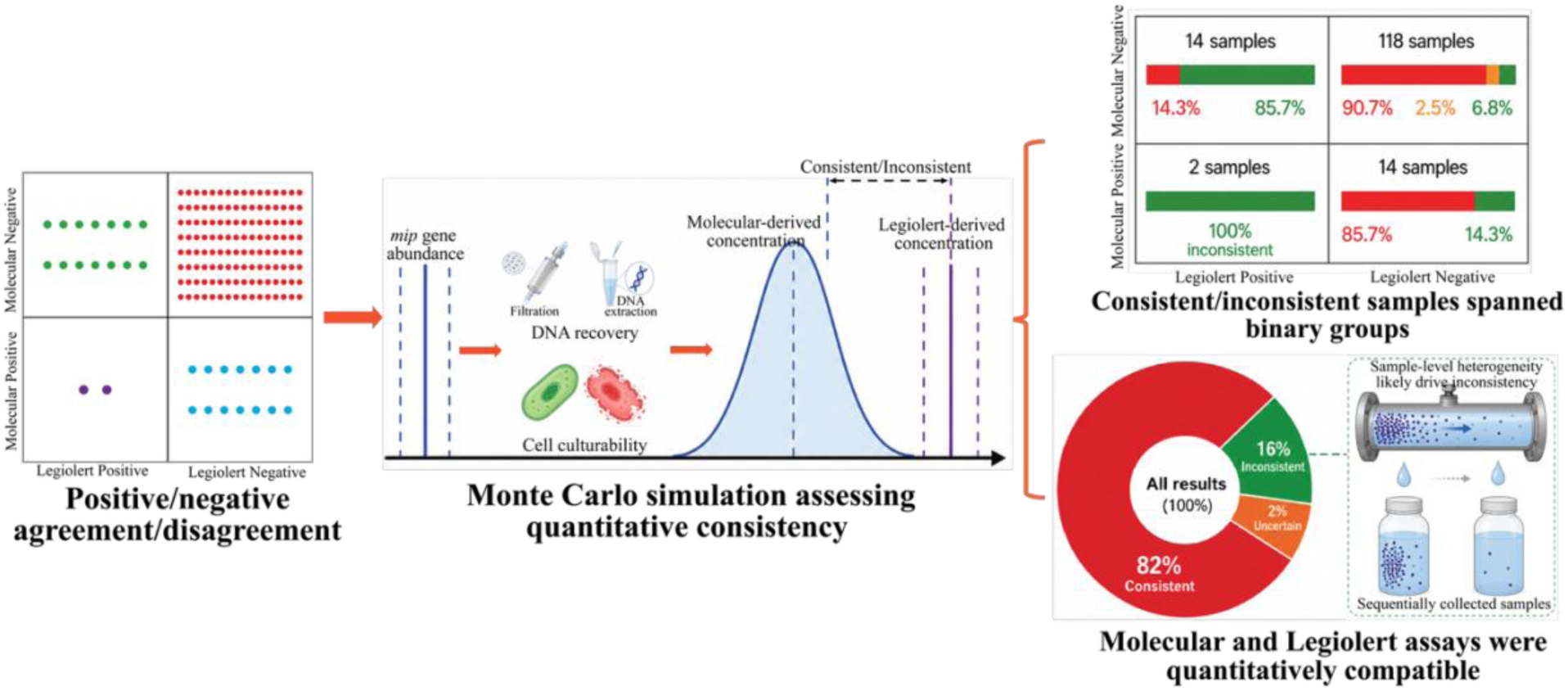

## Introduction

*Legionella spp.* are natural inhabitants of aquatic environments^1^ and are frequently detected in drinking water^1,2^. Although more than 60 *Legionella* species have been recognized^3^, *L. pneumophila* accounts for over 90% of clinical isolates associated with Legionnaires’ disease^4–6^, with serogroup 1 representing the predominant clinical serogroup^4,5^. However, the clinical dominance of *L. pneumophila* does not necessarily correspond to the environmental distribution of *Legionella* species and serogroups^1,5^. Drinking water systems can experience elevated temperatures, prolonged water age, stagnation, and residual disinfectant decay, favoring the regrowth of *L. pneumophila*^7–9^. This persistence is further promoted by biofilms and free-living amoebae, which provide nutrient-rich habitats and intracellular replication niches that protect *L. pneumophila* from environmental stressors and chemical disinfection^10^. Drinking water systems are recognized as important sources of *L. pneumophila* exposure^11–13^, with transmission occurring through inhalation of contaminated aerosols generated from potable-water fixtures, such as showers, or through aspiration of contaminated water, particularly in susceptible hospitalized populations^14–17^.

In the United States, routine *L. pneumophila* abundance monitoring is not federally mandated across public water systems^18,19^; however, when direct *L. pneumophila* testing is performed, standard plate culture-based method remains the regulatory reference approach^20,21^. These methods require 7 to 10 days of incubation and can be limited by poor recovery due to overgrowth of competing microorganisms^22,23^. Confirmatory testing of presumptive colonies requires specialized facilities and laboratory capacity^24^. These limitations can increase the risk of false-negative results and may provide unwarranted confidence in water-system risk management^22^. Alternative culture-based methods, such as the Legiolert assay, simplify sample processing and provide high specificity for *L. pneumophila*, but they still require 7 days of incubation^25–28^. These multi-day incubation requirements limit the usefulness of culture-based testing for rapid operational decision-making. Molecular assays, such as quantitative polymerase chain reaction (qPCR) and digital polymerase chain reaction (dPCR), provide a highly specific and rapid alternative for *L. pneumophila* detection^29^. However, molecular assays frequently yield higher detection frequencies and concentrations than culture-based methods, partly because they detect DNA from both culturable and non-culturable cells, as well as extracellular DNA^30–32^. The reverse scenario has also been documented, where samples yield positive results by culture-based assay but negative results by molecular assays^28,33,35^. This complicates interpretation of *L. pneumophila* monitoring results especially when these two assays lead to conflicting conclusions, which hinders the use of molecular assays as rapid complementary tools for culturable *L. pneumophila* monitoring. Previous studies have generally evaluated molecular-culture disagreement using direct assay-derived concentrations or binary positive/negative outcomes based on assay-specific thresholds^30–38^. Although incomplete DNA recovery and cell culturability are frequently discussed as possible explanations for disagreement, they have rarely been incorporated directly into quantitative consistency assessment. Consequently, it remains unclear whether molecular-culture disagreement reflects concentration-level incompatibility between assays or unaccounted methodological and physiological differences related to DNA recovery and cell culturability.

In this study, we developed a probabilistic Monte Carlo framework that incorporates sample-specific DNA recovery and cell culturability to evaluate quantitative consistency between molecular and Legiolert assays for *L. pneumophila* detection in source and finished drinking water. Specifically, we first characterized positive/negative agreement between the two assays, and assessed concentration-level compatibility between molecular-derived and Legiolert-derived culturable *L. pneumophila* concentrations with the Monte Carlo framework. By incorporating DNA recovery and cell culturability, this framework moves assay comparison beyond binary positive/negative agreement toward concentration-level compatibility and supports the use of molecular assays as rapid complementary tools for culturable *L. pneumophila* monitoring.

## Materials and Methods

### Sample collection and microbiological quantification

Detailed procedures for sample collection and microbiological quantification were described previously^39–41^. Briefly, water samples were collected from eight full-scale drinking water utilities, designated Utilities 2-9, located across diverse geographic regions of the United States. Samples included source water (n = 35), point-of-entry finished water (POE; n = 35), and drinking water distribution system samples (DWDS; n = 112). POE and DWDS samples were collectively defined as finished drinking water. *L. pneumophila* was quantified using the Legiolert® culture-based most-probable-number assay (Legiolert assay), with the Legiolert® Test and 100-mL Quanti-Tray®/Legiolert® trays (IDEXX Laboratories, Inc.), and a molecular assay targeting the *mip* gene^42^. The molecular assay was performed by dPCR using a QIAcuity™ Digital PCR System with QIAcuity Nanoplate 8.5k 96-well plates (QIAGEN, Hilden, Germany). Total cell counts (TCC) and intact cell counts (ICC) were quantified by flow cytometry, and heterotrophic plate counts (HPC) were quantified using the IDEXX HPC for Quanti-Tray® Test with 100-mL water samples (IDEXX Laboratories, Inc.). For each sampling event, molecular analysis and flow cytometry were performed in one laboratory on one sequentially collected water fraction, whereas HPC and Legiolert assay were performed in a second laboratory on a separate sample fraction.

### Positive/Negative Agreement and Threshold Adjustment for Molecular and Legiolert Assays

Positive/negative agreement between molecular and Legiolert assays was evaluated using assay-specific positivity thresholds. Molecular results were classified as positive when the measured *mip* gene concentration exceeded either half the limit of quantification (half-LOQ; 0.99 log10 gene copies per reaction) or the limit of blank (LOB; 0.77 log10 gene copies per reaction) depending on comparisons being considered (see Results and Discussion section). Legiolert results were classified as positive when the detected *L. pneumophila* concentration was at or above 1 MPN/100 mL. Each sample was assigned to one of four categories: molecular-positive/Legiolert-positive, molecular-positive/Legiolert-negative, molecular-negative/Legiolert-positive, or molecular-negative/Legiolert-negative.

Sample-specific DNA recovery and cell culturability were incorporated sequentially to convert the molecular positivity threshold into DNA-recovery-adjusted and culturability-adjusted thresholds. The sample-specific DNA recovery ratio was estimated as the fraction of theoretically present cellular DNA recovered after filtration and DNA extraction. Because the theoretical DNA mass depends on the assumed average genome size of the microbial cells in the sample, DNA mass per cell was first estimated from genome size as:

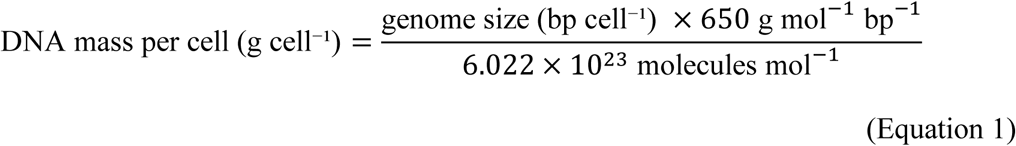

where 650 g mol⁻¹ bp⁻¹ was used as the average molecular weight of double-stranded DNA^43^, and 6.022 × 10^23^ molecules mol⁻¹ is Avogadro’s number. The median metagenome assembled genome (MAG)-derived genome size (3.07 Mbp) from the Drinking Water Genome Catalog was used as the representative genome size^44^.

The DNA recovery ratio was then calculated as:

### DNA mass extracted (g)

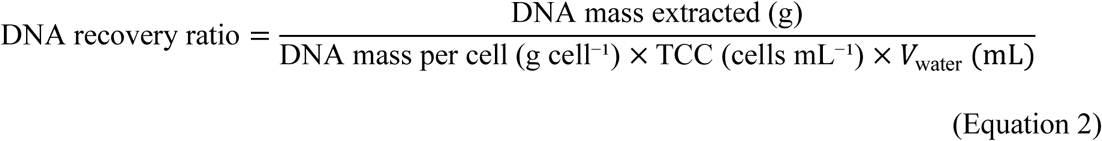

where TCC is the total cell count concentration measured by flow cytometry and *V*_water_ is the volume of water processed for DNA extraction.

The DNA-recovery-adjusted molecular threshold was calculated as:

### DNA-recovery-adjusted threshold (cells mL^−1^)

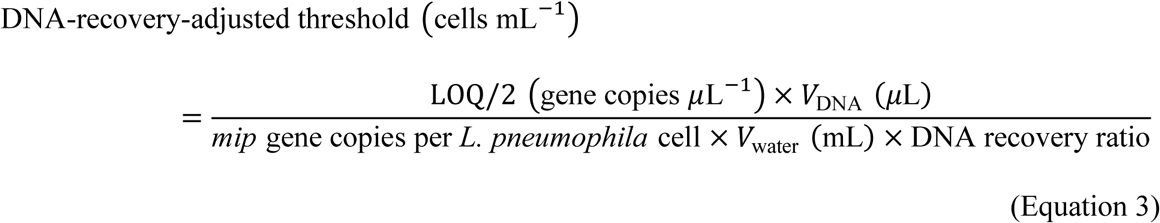

where LOQ/2 is the molecular positivity threshold used for threshold-adjusted interpretation,

*V*_DNA_ is the final DNA extract volume, and *V*_water_ is the volume of water processed for DNA extraction.

To further account for cell culturability, the DNA-recovery-adjusted threshold was converted into a culturability-adjusted threshold using a sample-specific bulk culturable ratio:

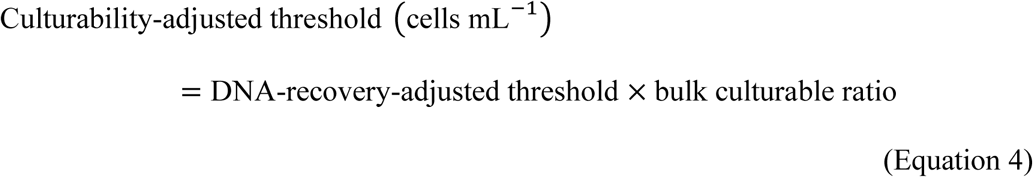

The bulk culturable ratio of the microbial community was approximated as:

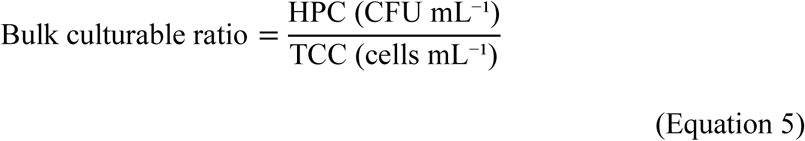

where HPC is the heterotrophic plate count. This ratio was used as proxy for the sample-specific culturability of *L. pneumophila*, because paired sample-specific quantification of total and culturable *L. pneumophila* cells was not available.

### Monte Carlo simulation to assess quantitative consistency between molecular and Legiolert assays

Monte Carlo simulation^45^ was used to estimate culturable *L. pneumophila* concentrations from molecular assay results and to evaluate quantitative consistency with Legiolert-derived concentrations. The framework converted measured *mip* gene concentrations to estimated culturable *L. pneumophila* concentrations by accounting for two factors: incomplete DNA recovery during sample processing and the culturability of *L. pneumophila*.

The estimated culturable *L. pneumophila* concentration was then calculated as:

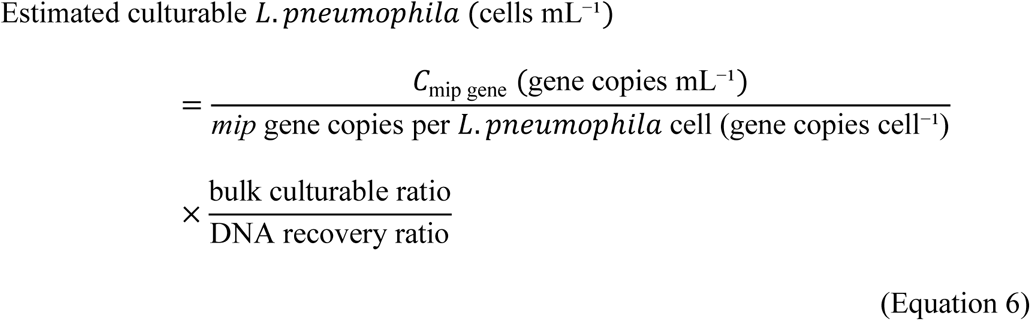

where *C*_mip_ is the *mip* gene concentration in the water sample. For molecular-positive samples, measured *mip* gene concentration was used directly in Equation 6. For molecular-negative samples, *mip* gene concentration was treated as left-censored below the molecular positivity threshold. Two censoring approaches were evaluated: uniform random draws between 0 and the threshold (uniform random-draw approach), and fixed substitution at the threshold (fixed upper-bound approach). The primary analysis used the half-LOQ^49,50^ threshold, and the uniform random-draw approach. Three MAG-derived genome-size assumptions from the Drinking Water Genome Catalog were evaluated in DNA recovery ratio calculation: Q1, median, and Q3, corresponding to 2.05, 3.07, and 4.25 Mbp, respectively^44^. The median genome size was used for the primary analysis, whereas Q1 and Q3 were used for sensitivity analyses. Samples missing TCC data were excluded because TCC was required to calculate both DNA recovery and bulk culturable ratios. Samples with calculated DNA recovery ratios greater than 1 or bulk culturable ratios greater than 1 were also excluded because these values were outside the physically interpretable range.

DNA recovery and bulk culturable ratio were incorporated into the Monte Carlo framework as sample-specific uncertain variables rather than fixed correction factors, because both ratios varied across samples and were estimated with measurement and biological uncertainty. To define appropriate uncertainty distributions, the empirical DNA recovery ratios and bulk culturable ratios were first fitted using candidate beta and logit-normal distributions. Model fit was evaluated using Akaike information criterion (AIC), Bayesian information criterion (BIC), Kolmogorov–Smirnov tests, and Q–Q plots. The logit-normal distribution provided the better fit and was therefore used to define the overall distributional form for both ratios. To preserve between-sample differences while incorporating uncertainty around each sample-specific estimate, local uncertainty distributions were centered on each sample’s observed DNA recovery ratio and bulk culturable ratio. Local uncertainty was modeled on the logit scale, with the local standard deviation defined as a fraction of the corresponding global logit-scale standard deviation^46–48^. Local standard deviation fractions of 5–25% were tested for both ratios to evaluate sensitivity to the assumed uncertainty magnitude.

For each sample, 10,000 Monte Carlo iterations were performed. In each iteration, DNA recovery ratio, bulk culturable ratio, and, when applicable, censored *mip* gene concentrations were sampled to estimate culturable *L. pneumophila* concentration. The resulting simulated values were used to calculate the probability of quantitative consistency with the Legiolert assay. For Legiolert-negative samples, consistency was defined as the probability that the simulated culturable *L. pneumophila* concentration was below the Legiolert detection threshold (0.01 MPN/mL). For Legiolert-positive samples, consistency was defined as the probability that the simulated concentration was within a numerical tolerance (0.005 MPN/mL) of the Legiolert-derived concentration. Samples were classified as consistent, uncertain, or inconsistent when the probability of consistency was ≥0.7, 0.3-0.7, or <0.3, respectively. The primary Monte Carlo analysis used the median genome-size assumption, logit-normal uncertainty distributions, uniform random draws for molecular-negative samples, and a local standard deviation equal to 5% of the global logit-scale standard deviation. Sensitivity analyses included testing genome-size assumptions, censoring approaches, and local uncertainty magnitudes.

### *Post hoc* comparison of bulk microorganisms and *L. pneumophila*-specific culturable ratios

A *post hoc* comparison was conducted to evaluate whether the bulk culturable ratio adequately represented *L. pneumophila*-specific culturability. Molecularly estimated total *L. pneumophila* concentration was calculated from the measured *mip* gene concentration after correction for *mip* gene copy number per cell and DNA recovery:

### Molecularly estimated total *L*. *pneumop*ℎ*ila* (cells mL⁻¹)

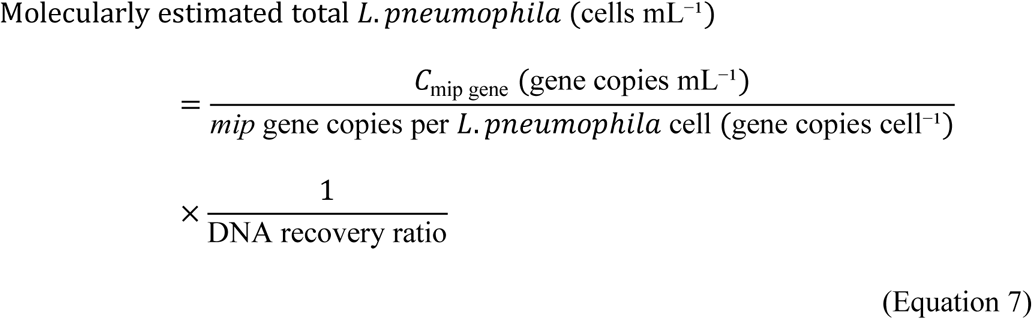

The *L. pneumophila*-specific culturable ratio was calculated as:

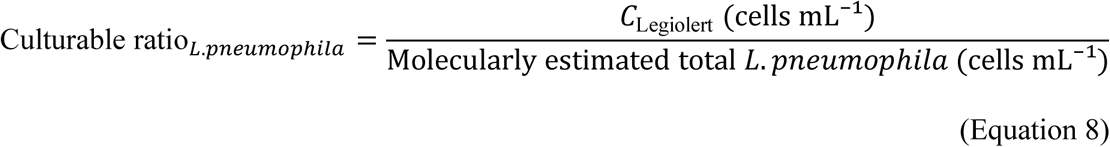

where *C*_Legiolert_is the Legiolert-derived *L. pneumophila* concentration, and *C*_mip gene_is the *mip* gene concentration. For molecular-negative samples, *C*_mip gene_was substituted with half-LOQ.

The paired difference between the *L. pneumophila*-specific and bulk culturable ratio was calculated as:

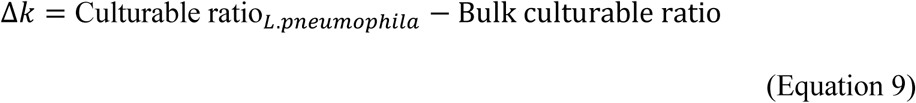

This comparison was used only as a post hoc diagnostic analysis and the *L. pneumophila*-specific culturable ratio was derived using the Legiolert assay and therefore was not used as an input parameter in the Monte Carlo simulation.

## Statistical analysis

McNemar’s exact test was used to evaluate asymmetry between molecular-positive/Legiolert-negative and molecular-negative/Legiolert-positive disagreement. Paired Wilcoxon signed-rank tests were used for paired comparisons between Legiolert-derived and Monte Carlo-estimated culturable *L. pneumophila* concentrations and between HPC and ICC. Wilcoxon rank-sum tests were used to compare physicochemical variables between consistent and inconsistent samples. Principal component analysis (PCA) and permutational multivariate analysis of variance (PERMANOVA) based on Euclidean distance matrices were used to assess multivariate differences in water chemistry between consistency groups. Statistical significance was defined as P < 0.05.

## Results and Discussion

### Positive/negative disagreement was not fully resolved by DNA-recovery-and culturability-adjusted thresholds

Using half-LOQ as the molecular positivity threshold, 11.2% of samples were classified as molecular-positive for *L. pneumophila*, which was slightly lower than the positivity rate obtained with the Legiolert assay (12.3%; **Figure 1A**). At the sample level, 21.3% of samples showed positive/negative disagreement between the two assays. Specifically, 10.1% of samples were molecular-positive but Legiolert-negative, whereas 11.2% were molecular-negative but Legiolert-positive. Only 1.1% of samples were positive by both assays, while 77.7% were negative by both assays. McNemar’s exact test showed no significant asymmetry between the two disagreement patterns (P > 0.05), indicating that the two discordant patterns occurred at comparable frequencies. When the Legiolert assay was used as the reference method, based on its previously demonstrated statistical equivalence with the standard culture method^36^, the molecular assay showed low sensitivity (9.1%) and positive predictive value (PPV; 10.0%), but relatively high specificity (88.5%) and negative predictive value (NPV; 87.4%). Thus, the relatively high overall accuracy (78.8%; **Figure 1C**) was driven primarily by both-negative samples, whereas positive agreement between the two assays remained limited.

**Figure 1.**
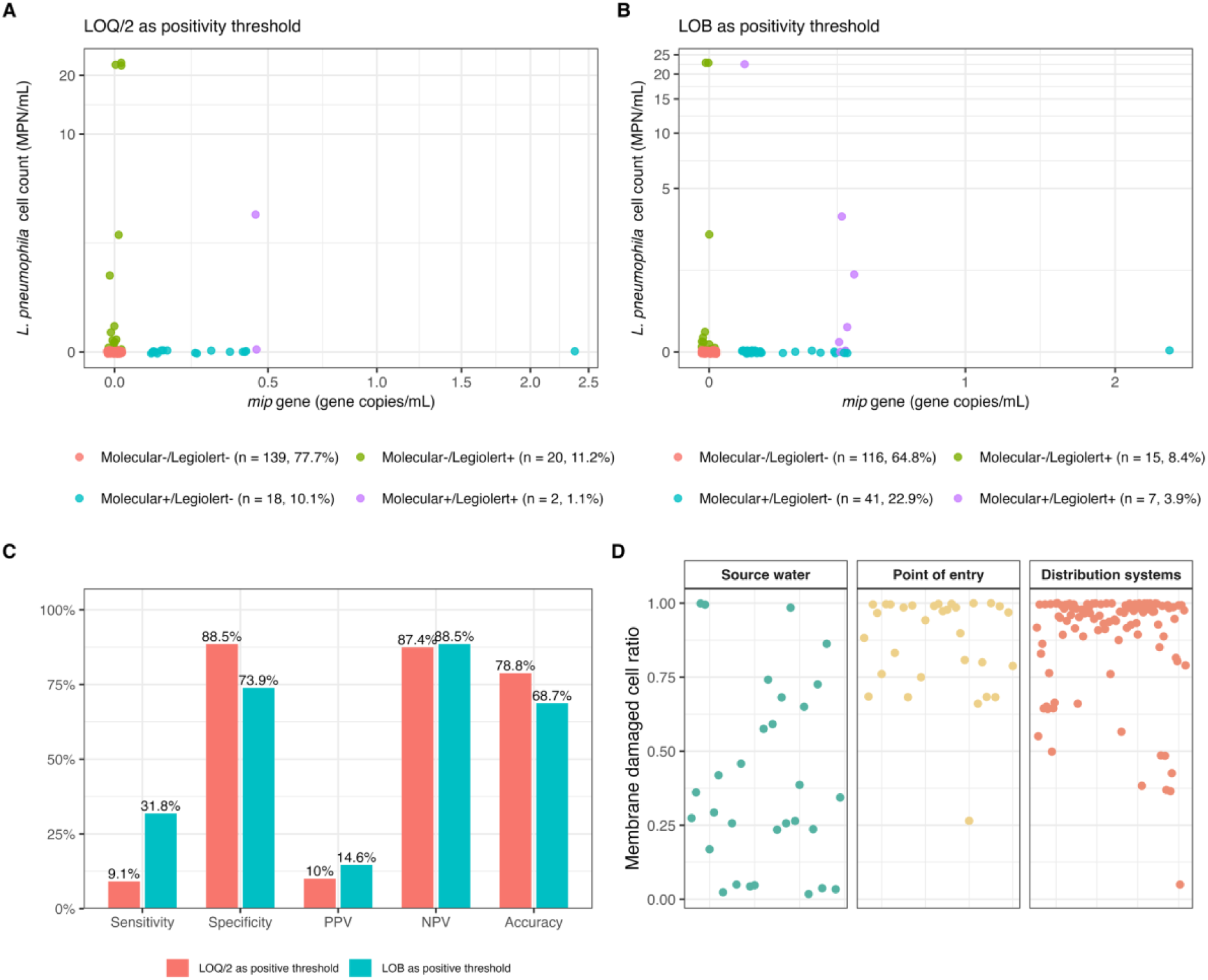
Sample-level comparison of positive/negative detection of *Legionella pneumophila* between molecular and Legiolert assays using (A) half-LOQ and (B) LOB as the molecular positivity thresholds. (C) Diagnostic performance of the molecular assay under each threshold, including sensitivity, specificity, positive predictive value (PPV), negative predictive value (NPV), and accuracy, with Legiolert positive/negative detection as the reference. (D) Membrane-damaged cell ratio in water samples across source water, point of entry, and distribution system.

LOB was applied as a more permissive threshold to assess whether low sensitivity resulted from an overly stringent molecular positivity cutoff. Under the LOB threshold, the number of molecular-positive samples increased (**Figure 1B**), and molecular assay sensitivity increased to 31.8%, while specificity and accuracy decreased to 73.9% and 68.7%, respectively (**Figure 1C**). However, positive agreement between the two assays remained limited: only seven samples were positive by both assays (3.9%), whereas 116 samples were negative by both assays (64.8%). Consistent with this pattern, the PPV remained low (14.6%), while the NPV remained high (88.5%). McNemar’s exact test showed significant asymmetry in disagreement patterns (P < 0.05), consistent with the higher proportion of molecular-positive/Legiolert-negative samples. Thus, although lowering the molecular positivity threshold improved sensitivity, this improvement occurred at the expense of specificity and did not substantially improve overall accuracy between the molecular and Legiolert assays. These results indicate that positive/negative disagreement was not explained solely by the molecular positivity threshold.

Disagreement between qPCR and standard plate-culture method for *L. pneumophila* detection has been widely reported^31,32,37^. A seven-study meta-analysis demonstrated equal or higher positive-sample proportions detected by qPCR than culture, with differences ranging from 0 to 63%^32^. When the standard plate-culture method was used as the reference, qPCR frequently showed high sensitivity but low specificity^30,31^. For example, one study of *L. pneumophila* detection in drinking water from hospitals and hotels reported 100% sensitivity (32/32) but only 44.2% specificity (19/43)^31^. However, molecular assay showed low sensitivity but relatively high specificity relative to Legiolert assay in this study. This opposite sensitivity-specificity pattern was unlikely to result simply from replacing standard plate-culture with Legiolert assay as the culture-based comparator because Legiolert assay has been shown to be statistically equivalent to the standard plate-culture method for *L. pneumophila* detection^25^. In addition, a previous study comparing viability qPCR with both Legiolert and the standard plate-culture method reported similar sensitivity-specificity patterns across both culture-based methods^37^. False-positive Legiolert results also were unlikely to be the primary explanation for the molecular-negative/Legiolert-positive pattern. Although false-positive Legiolert results cannot be excluded for individual disagreed samples, previous evaluations of the Legiolert assay have typically reported low false positive rates of approximately 2.1-4.8%^25–27^. Even a drinking water study reporting a higher variance found only 33.3% false-positive rate^51^. In contrast, attributing all 20 molecular-negative/Legiolert-positive samples in the present study to Legiolert false positives results in an implausibly high 90.9% false-positive rate across all Legiolert-positive results.

The observed positive/negative disagreement was therefore evaluated by sequentially adjusting the molecular threshold for two assay-relevant factors: incomplete DNA recovery during water filtration and DNA extraction, and limited culturability of *L. pneumophila*. After accounting for incomplete DNA recovery alone, the DNA-recovery-adjusted threshold was higher than the Legiolert threshold across samples (590 ± 1304 cells/100 mL *vs.* 1 MPN/100 mL). This threshold difference indicates that samples containing culturable *L. pneumophila* above the Legiolert threshold but below the DNA-recovery-adjusted threshold could be classified as Legiolert-positive but molecular-negative. The sample-specific bulk culturable ratio was then incorporated to convert the DNA-recovery-adjusted threshold into culturability-adjusted threshold. This adjustment was particularly relevant for molecular-positive/Legiolert-negative samples because molecular detection can capture *L. pneumophila mip* genes from non-culturable or poorly culturable fractions, including viable-but-nonculturable cells^54–56^, membrane-damaged cells^57,58^, and cells residing within free-living amoebae^33^, and extracellular DNA^59,60^. For molecular-positive/Legiolert-negative samples, this threshold was lower than the Legiolert threshold in 15 of 18 samples, supporting limited *L. pneumophila* culturability as a major explanation for this disagreement pattern. This interpretation was consistent with the high fraction of membrane-damaged cells in the bulk microbial community, with fractions averaging 0.38 ± 0.29 in source water, 0.87 ± 0.17 in POE samples, and 0.88 ± 0.19 in DWDS samples (**Figure 1D**). Although this metric was not *L. pneumophila*-specific, it suggests that membrane-damaged cells were common in these samples and could contribute to molecular-positive/Legiolert-negative samples. However, the same correction explained only a minority of molecular-negative/Legiolert-positive samples: the culturability-adjusted threshold exceeded the Legiolert threshold in only 4 of 20 samples. Thus, threshold mismatch after accounting for DNA recovery and cell culturability could explain only part of the Legiolert-positive/molecular-negative disagreement, highlighting the limitation of threshold-based classification. The remaining discordant samples likely involved confounding factors, such as heterogeneity between sequential samples analyzed by the two assays, uncertainty in the sample-specific DNA recovery and bulk culturable ratios, or occasional false-positive Legiolert reactions. Therefore, the analysis was extended beyond positive/negative agreement to evaluate quantitative consistency between molecular-and Legiolert-derived culturable *L. pneumophila* concentrations.

### Quantitative consistency classifications were robust across Monte Carlo parameterizations

Monte Carlo simulation incorporated DNA recovery and cell culturability into the quantitative comparison between molecular and Legiolert assays. Because genome size was required to estimate culturable *L. pneumophila* cell concentrations from *mip* gene copies, 1,141 MAGs from previous drinking water studies were used to approximate the plausible genome-size distribution^44^. The sizes of these MAGs ranged from 0.42 to 10.56 Mbp, with Q1, median, and Q3 values of 2.05, 3.07, and 4.25 Mbp, respectively (**Figure 2A**). Beta and logit-normal distributions were fitted to the DNA recovery ratio and bulk culturable ratio. Using the median MAG-derived genome size, the logit-normal model provided a better fit for the DNA recovery ratio than the beta model, as indicated by lower AIC and BIC values (−262.93 vs −238.48 and −256.94 vs −232.49, respectively; **Figures 2B and S1**). The Kolmogorov–Smirnov tests for the logit-normal model was not significant (P = 0.714), indicating that this model adequately captured the empirical distribution of DNA recovery ratios. For the bulk culturable ratio, the logit-normal model also had lower AIC and BIC values than the beta model (**Figures 2C and S2**), although the Kolmogorov–Smirnov tests remained significant (P = 8.90 × 10⁻⁵). This deviation was mainly associated with a few samples with bulk culturable ratios greater than 5%, up to 15.8% (**Figures 2D**), indicating greater heterogeneity in cell culturability across source water and finished drinking water. When the fitting procedure was repeated using the Q1 and Q3 MAG-derived genome sizes, the same model-selection pattern was observed for both DNA recovery and bulk culturable ratios (**Figures S3–S6**), supporting the robustness of the logit-normal approximation across the evaluated genome-size assumptions.

**Figure 2.**
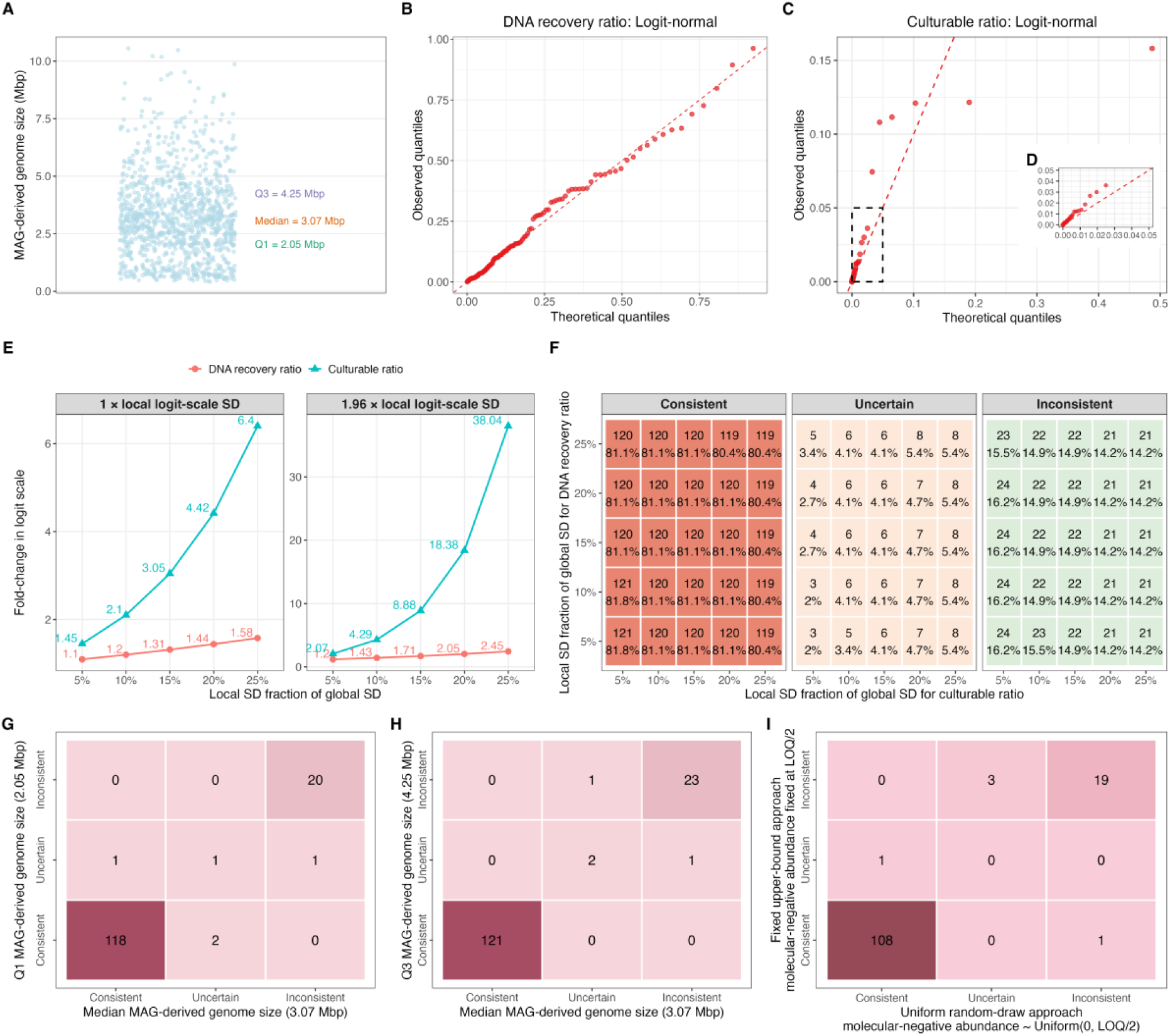
(A) Distribution of MAG-derived genome sizes retrieved from previous drinking water studies. Monte Carlo parameterization and sensitivity analysis using the median MAG-derived genome size of 3.07 Mbp. (B, C) Q–Q plots of the logit-normal fits for the DNA recovery ratio (B) and bulk culturable ratio (C); the inset in (C) shows the low-ratio region (D). (E) Variation in logit-scale corresponding to 1 × and 1.96 × local logit-scale standard deviation for the DNA recovery ratio and bulk culturable ratio across local standard deviation fractions. (F) Consistency classifications across combinations of local standard deviation fractions of DNA recovery and bulk culturable ratio. (G, H) Sample-level comparison of consistency classifications using the median MAG-derived genome-size versus the Q1 (G) and Q3 (H). (I) Sample-level comparison of consistency classifications for molecular-negative samples using the fixed upper-bound approach and the uniform random-draw approach.

The logit-normal model was then used to define the local uncertainty structure for both parameters. For each sample, the DNA recovery ratio and bulk culturable ratio were centered on their sample-specific values, while the local standard deviation was set to 5-25% of the corresponding global logit-scale standard deviation. For DNA recovery ratio, increasing the local standard deviation produced relatively modest variation, with fold-changes of 1.10-1.58 for the 1 × local logit-scale standard deviation interval and 1.20-2.45 for the 1.96 × local logit-scale standard deviation interval (**Figure 2E**). In contrast, the bulk culturable ratio showed much broader variation, with fold-changes of 1.45-6.40 and 2.07-38.04 for the corresponding intervals. However, the consistency classification was stable across local standard deviation assumptions (**Figure 2F**). Across all 25 combinations of local standard deviation fractions of DNA recovery ratio and bulk culturable ratio, 119-121 samples were classified as consistent, 3-8 as uncertain, and 21-24 as inconsistent. Increasing bulk culturable ratio uncertainty mainly shifted a small number of samples for the consistency classifications. When the analysis was repeated using the Q1 and Q3 MAG-derived genome sizes, the overall variation patterns and consistency classifications remained similar (**Figures S7 and S8**). In addition, most samples retained the same consistency classification when the assumed genome size was changed from the median MAG-derived value to the Q1 or Q3 value, with only slightly shifts between consistent and uncertain or between uncertain and inconsistent (**Figures 2G, H**). Similarly, replacing the uniform random-draw approach with the fixed upper-bound approach caused limited reclassification for molecular-negative samples (**Figure 2I**). Collectively, these sensitivity analyses showed that most samples retained the same consistency classifications across the genome-size assumptions, local standard deviation fractions selected, and censoring approaches for molecular-negative samples. Therefore, the final Monte Carlo simulation used the median MAG-derived genome size, the uniform random-draw approach for molecular-negative samples, and a local standard deviation equal to 5% of the corresponding global logit-scale standard deviation.

### Molecular and Legiolert assays were quantitatively compatible for culturable *L. pneumophila* concentrations

Across the final Monte Carlo analysis, 121 of 148 samples were classified as quantitatively consistent in culturable *L. pneumophila* concentration, whereas 24 were inconsistent and only 3 were uncertain (**Figure 3A**). This high proportion of consistent samples indicates that molecular and Legiolert assays were generally quantitatively compatible for culturable *L. pneumophila* concentrations. Positive/negative agreement was then compared with quantitative consistency (**Figure 3B**). Among the both-negative samples, 107 of 118 were quantitatively consistent, whereas 3 were uncertain and 8 were inconsistent. Among the molecular-positive/Legiolert-negative samples, 12 of 14 were quantitatively consistent and 2 were inconsistent. In contrast, the two both-positive samples were quantitatively inconsistent, and 12 of 14 molecular-negative/Legiolert-positive samples were quantitatively inconsistent. Although both-negative agreement was largely associated with quantitative consistency, positive agreement did not necessarily indicate quantitative consistency, and disagreement did not necessarily indicate quantitative incompatibility. Thus, quantitatively consistent and inconsistent samples were distributed across both binary agreement and disagreement classifications, demonstrating that positive/negative classifications did not reliably capture quantitative consistency in culturable *L. pneumophila* concentration. These results highlight the need of quantitative consistency evaluation by incorporating sample-specific DNA recovery and cell culturability. In addition to the extensively reported positive/negative disagreement, previous studies have also compared *L. pneumophila* concentrations derived from molecular and culture-based assays. These studies generally relied on direct comparisons of assay-derived concentrations and reported variable correlations between the two assays^33,34^. Molecular assays generally yielded higher estimated concentrations than culture-based assays, with reported molecular-to-culture concentration ratios ranging from approximately 1:1 to 100:1^32,34,38^. However, this trend was not consistent across all samples, and some studies also reported samples in which culture-based concentrations exceeded molecularly derived concentrations^28,33,35^. These variable differences indicate that the relationship between molecular-derived and culture-based *L. pneumophila* concentrations was sample specific rather than governed by a consistent proportion. Therefore, direct interpretation of assay-derived concentrations without accounting for sample specific DNA recovery and cell culturability may misrepresent the quantitative relationship between molecular and culture-based assays. The Monte Carlo framework addresses this limitation by incorporating these two sample-specific factors and propagating their uncertainty in quantitative comparison between molecular and Legiolert assays.

**Figure 3.**
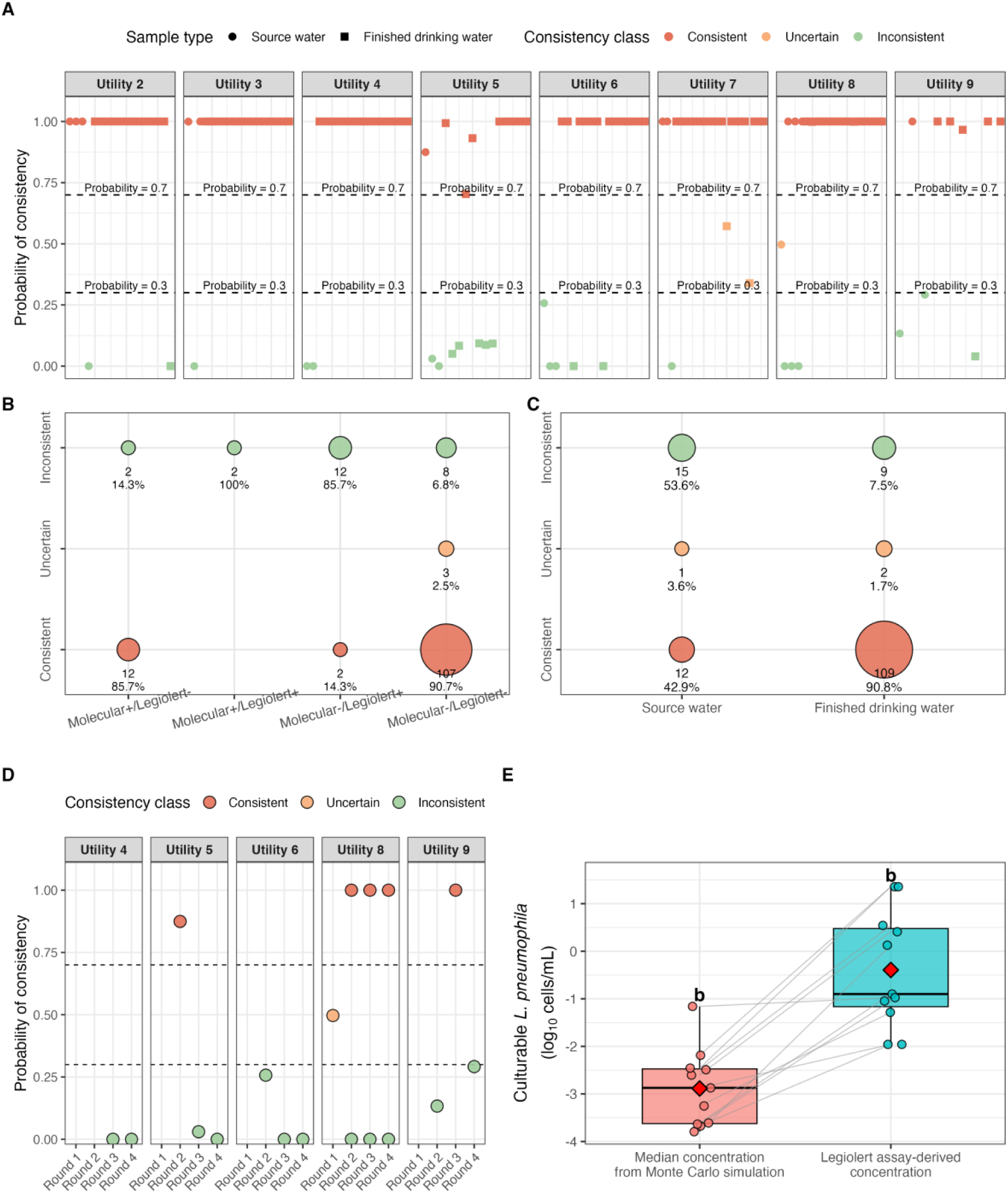
(A) Sample-level probability of consistency between molecular-and Legiolert-derived culturable *L. pneumophila* concentrations across utilities (B) Distribution of quantitative consistency classifications in positive/negative classifications (C) Distribution of quantitative consistency classifications in source water and finished drinking water. (D) Source water samples across sampling rounds for utilities with repeated inconsistent classifications. (E) Median molecular-derived culturable *L. pneumophila* concentration estimated from Monte Carlo simulation, and the Legiolert-derived concentration among inconsistent source water samples.

Among source water samples, 53.6% (15/28) were quantitatively inconsistent, which accounted for 62.5% of all inconsistent samples (**Figure 3C**). Of these 15 inconsistent source water samples, 9 were molecular-negative/Legiolert-positive, 2 were molecular-positive/Legiolert-negative, 2 were negative by both assays, and 2 were positive by both assays. The inconsistency recurred at several specific sampling sites across sampling rounds, accounting for 80% of inconsistent source water samples (**Figure 3D**). Specifically, source water samples from Utility 4 were classified as inconsistent in Rounds 3 and 4; Utility 5 in Rounds 3 and 4; Utility 6 in Rounds 2, 3, and 4; Utility 8 in Rounds 2, 3, and 4; and Utility 9 in Rounds 2 and 4. This quantitative inconsistency may result from within-sample heterogeneity in source water^61^, particularly because separate sample aliquots were used for these two assays in this study. Previous research showed that within-sample variability accounted for 62-100% of the total variability in measured surface-water chemistry variables^61^. For inconsistent source water samples, the Legiolert assay derived culturable *L. pneumophila* concentration was significantly higher than the median concentration estimated from the Monte Carlo simulation (-0.39 ± 1.21 vs-2.89 ± 0.81 log10 cells/mL, P < 0.05; **Figure 3E**). In the most extreme case, the Legiolert-derived concentration was 22.7 cells/mL, which was 11,350-fold higher than the median molecular-derived concentrations (0.002 cells/mL). Together, the large differences between molecular-and Legiolert-derived culturable *L. pneumophila* concentrations and the recurrence of inconsistency at the same source-water sites suggested that source-water heterogeneity likely contributed to the observed quantitative inconsistency. Previous studies have reported false-positive or cross-reactive Legiolert reactions associated with non-target bacteria, including *Brevundimonas vesicularis*, *Brevundimonas diminuta*, *Sphingomonas koreensis*, *Ochrobactrum intermedium*, and *Elizabethkingia anophelis* in drinking water^62^, as well as *Proteus mirabilis* and *Serratia marcescens* in laboratory cross-reactivity testing with bacterial isolates^63^. Although the present framework could not rule out false-positive Legiolert reactions for individual samples, attributing all quantitative inconsistencies to false-positive Legiolert results would not be justified, given the high Legiolert specificity from previous validation studies^25–27^.

Only 7.5% (9/120) of finished drinking water samples were quantitatively inconsistent, including three molecular-negative/Legiolert-positive samples and six both-negative samples (**Figure 3C**). This reduced rate compared with source water suggests that quantitative inconsistency decreased after treatment and distribution, potentially reflecting reduced within-sample heterogeneity due in part to greater hydraulic mixing in distribution systems than source water^64–66^. Drinking water in distribution systems typically experiences turbulent pipe-flow conditions during normal operation, particularly during periods of active demand or flushing^66^. In contrast, source waters may be more susceptible to within-sample heterogeneity because partial microbial biomass was associated with suspended particles or sediment-derived material^67,68^, which are not uniformly distributed in source water. Nevertheless, quantitative inconsistency was not randomly distributed but repeated at specific sampling sites, including sites 5-2 and 5-3, which were classified as inconsistent in both Rounds 2 and 3 (**Figure 4A**). These repeated cases accounted for 5 of the 9 inconsistent finished drinking water samples, suggesting that localized site conditions may have contributed to quantitative inconsistency. Additional atypical microbial indicators further supported this possibility. For example, HPC concentrations were significantly higher than ICC in these five repeated inconsistent samples and in site 9-2 in Round 3 (n = 6 samples; 3.34 ± 0.08 log10 CFU/mL vs 2.80 ± 0.49 log10 cells/mL, P < 0.05; **Figure 4B**). To minimize confounding by source water differences, the bulk culturable ratio of inconsistent finished drinking water samples was compared with that of consistent samples from the same utility and sampling round. The bulk culturable ratio was 1.5-, 24.15-, and 54.91-fold higher in inconsistent samples from Utility 5 in Round 2, Utility 5 in Round 3, and Utility 9 in Round 3, respectively (**Figure 4C**). In addition, the Legiolert-derived culturable *L. pneumophila* concentration was higher than HPC concentration in inconsistent finished drinking water sample from site 2-1 in Round 4 (22.72 *L. pneumophila* cells/mL vs 0 CFU/mL). Although HPC, flow cytometry, and Legiolert assay target different microbial fractions and lack direct comparability, cross-indicator discrepancies provide circumstantial evidence that within-sample heterogeneity underlies the inconsistencies observed in finished drinking water. Together, repeated site-level inconsistency, elevated bulk culturable ratios in inconsistent samples, and anomalously high Legiolert-derived culturable *L. pneumophila* concentrations collectively suggest that localized within-sample heterogeneity likely contributed to quantitative inconsistency at specific finished-drinking-water sampling sites. Previous studies support the plausibility of within-sample heterogeneity during water collection. HPC, ICC, and TCC have been shown to vary across sequentially collected water fractions, with overall declines across the first 0-5 L of collected water but additional fluctuations among fractions^69^. Similar temporal fluctuations were also reported when bacterial dynamics were evaluated over flushing time^70^. These findings indicate that water collected sequentially from the same outlet may not be compositionally uniform over the sampling period. Hydraulic conditions provide a plausible mechanism for this heterogeneity. Previous studies have demonstrated that laminar flow can occur in dead ends of distribution systems and in premise plumbing because of small pipe diameters or lower velocities^71–73^, which can limit hydraulic mixing and generate time-varying water quality at the sampling outlet. Tracer-transport experiments further showed that steady turbulent flow produced well-mixed, uniform tracer concentrations, whereas laminar and accelerating flows produced spatially nonuniform or temporally multi-peaked concentration profiles because of limited mixing and longitudinal differential advection^65^. Therefore, the molecular and Legiolert assays likely captured different microbial fractions, contributing to quantitative inconsistency. These findings highlight the need for standardized *L. pneumophila* sampling procedures that define collection, mixing, aliquot-splitting, and replication strategies to reduce within-sample heterogeneity, particularly when multiple parameters are measured for cross-assay comparisons and modeling. The appropriate sampling design will depend on the spatial and temporal distribution of *L. pneumophila* within the water sample, especially when cells are particle-associated or intermittently released from biofilms and plumbing deposits. Incorporating the statistical sampling components would reduce within-sample bias, improve quantitative comparability across assays, and strengthen large-scale risk assessments of *L. pneumophila* in drinking water. False-positive or cross-reactive Legiolert reactions may also have contributed to the Legiolert-positive inconsistent finished drinking water samples. For example, site 6-3 was inconsistent in both Rounds 2 and 3 despite normal microbial indicator levels and bulk culturable ratios. This pattern suggests that sample-level heterogeneity was less evident in these two samples and occasional false-positive or cross-reactive Legiolert reactions may have contributed to the observed inconsistency.

**Figure 4.**
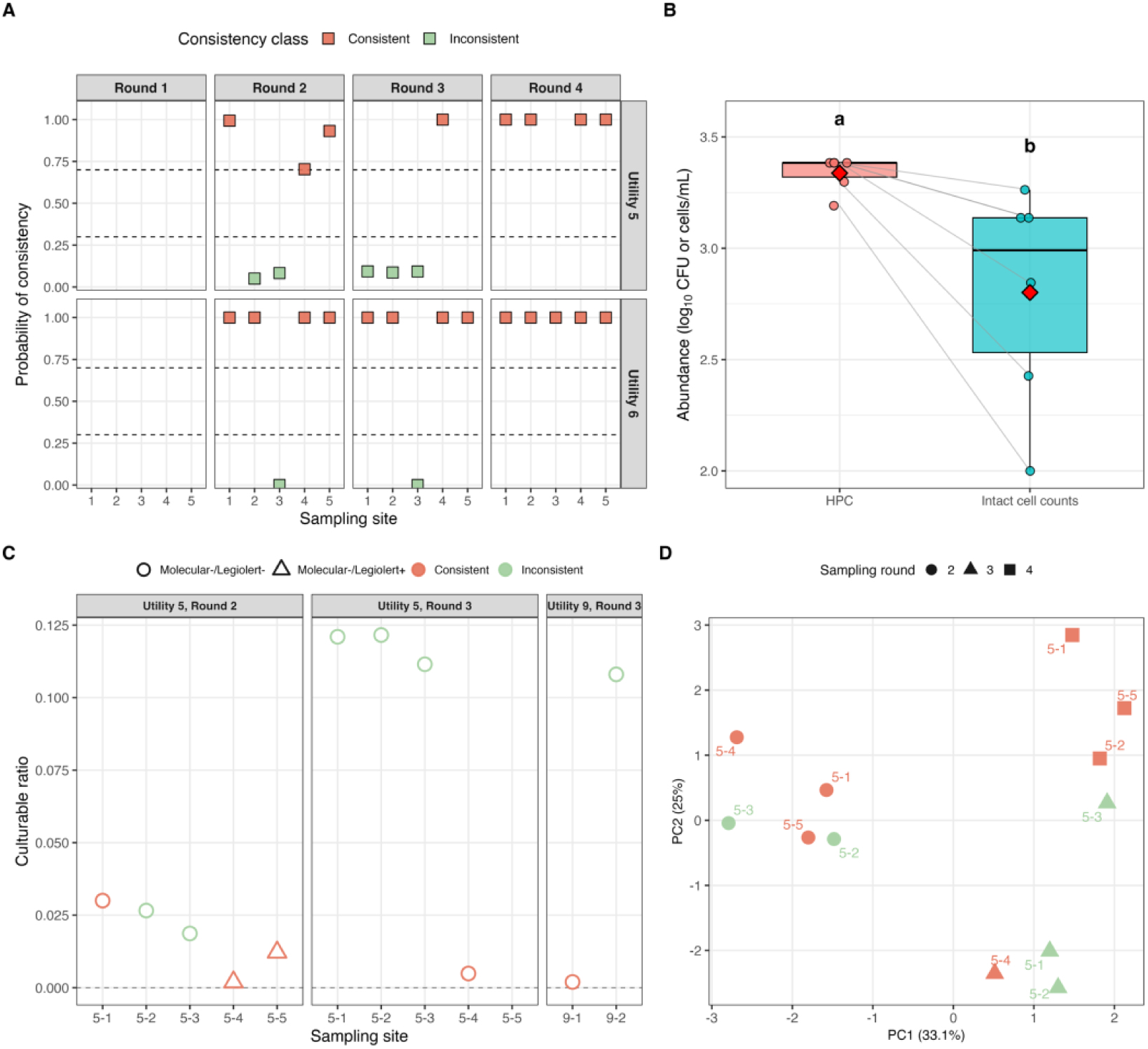
(A) Probability of consistency for finished drinking water samples from Utilities 5 and 6 across sampling sites and rounds. (B) Heterotrophic plate counts and intact cell counts in inconsistent finished drinking water samples from sites 5-2 and 5-3 in Round 2 and sites 5-1, 5-2, 5-3, and 9-2 in Round 3; Red diamonds indicate mean values. Different lowercase letters indicate significant differences (P < 0.05). (C) Bulk culturable ratio in selected finished drinking water samples from Utility 5 in Rounds 2 and 3 and Utility 9 in Round 3. (D) Principal component analysis of measured physicochemical parameters in Utility 5 finished drinking water samples.

Another possible explanation for the quantitative inconsistency is that the bulk culturable ratio may not accurately represent *L. pneumophila*-specific culturability^74,75^. HPC reflects the culturable fraction of the broader heterotrophic community, whereas *L. pneumophila* culturability may be governed by organism-specific responses to disinfectant exposure and other water-quality stresses^76–79^. To examine whether this quantitative inconsistency is associated with consistent shift in water chemistry, water chemistry profiles were compared between consistent and inconsistent finished drinking water samples in Utility 5. PCA showed that the overall water chemistry profile was more strongly structured by sampling rounds than the consistency classifications. Consistent and inconsistent samples were not clearly separated, as supported by PERMANOVA (P > 0.05; **Figure 4D**). To reduce potential confounding from sampling rounds and maintain a more balanced comparison between consistency classifications, univariate statistical analyses were performed using only Utility 5 finished drinking water samples from Rounds 2 and 3. No significant differences were observed between consistent and inconsistent samples for the measured physicochemical variables, including pH, temperature, ammonia, DOC, sulfate, nitrate, Fe, Cu, Zn, Mg, and total chlorine (P > 0.05; **Figure S9**). These results indicated that the quantitative inconsistency was not driven by systematic shifts in the measured physicochemical parameters.

### Bulk culturable ratio broadly approximated *L. pneumophila*-specific culturability

A *post hoc* diagnostic comparison was conducted to evaluate whether the bulk culturable ratio adequately represented *L. pneumophila*-specific culturability. In source water, bulk culturable ratio of consistent samples was higher than or equal to the *L. pneumophila*-specific culturable ratio, with differences ranging from-1.37% to 0% and an average difference of-0.20% (**Figure 5A**). Inconsistent source water samples showed more variable deviations. Ten inconsistent source water samples showed relatively small differences, ranging from-7.44% to 4.48%, whereas the remaining five inconsistent samples showed much higher *L. pneumophila*-specific culturable ratios, with differences ranging from 101.3% to 729.9%. A similar pattern was observed in finished drinking water (**Figure 5B**). Bulk culturable ratio of consistent finished drinking water samples was also higher than or equal to the *L. pneumophila*-specific culturable ratios, with differences ranging from-3.00% to 0% and an average difference of-0.10%. Among inconsistent finished drinking water samples, four showed relatively small differences, ranging from-2.66% to 0.53%. The remaining five samples showed larger deviations, including four samples with differences ranging from-10.8% to-12.2%, and one sample with an extreme difference of 20,425%. The close agreement between the bulk and *L. pneumophila*-specific culturable ratio in most samples, particularly consistent samples, supports the use of bulk culturable ratio as a reasonable proxy for *L. pneumophila* culturability in the Monte Carlo framework.

**Figure 5.**
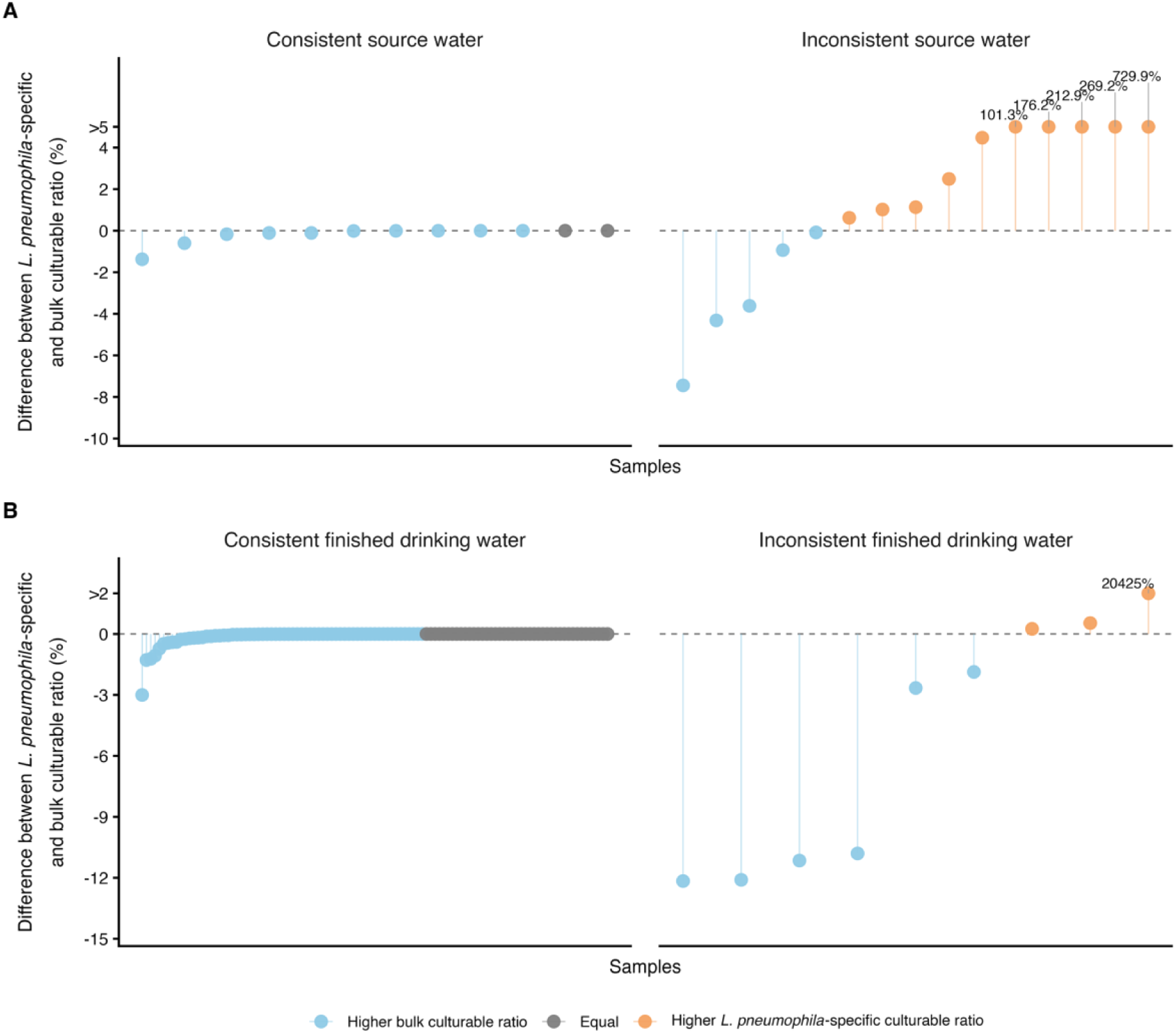
*Post hoc* comparison of *L. pneumophila*-specific culturable ratio and bulk culturable ratio in (A) source water and (B) finished drinking water samples.

### Environmental Implications

Persistent discrepancies between molecular and culture-based *L. pneumophila* detection have impeded the standardization of monitoring protocols and delayed the regulatory adoption of rapid molecular assays. This study directly challenges that paradigm by demonstrating that once sample-specific cell culturability and DNA recovery are accounted for, molecular-derived *L. pneumophila* concentrations can align quantitatively with Legiolert-derived concentrations. These results strongly support the deployment of molecular assays as a rapid, routine monitoring approach. When coupled with robust uncertainty estimates, rapid molecular screening could enable timely and more proactive operational interventions, such as targeted flushing and disinfectant adjustment, to promptly mitigate *L. pneumophila* proliferation and associated health risks. Although the current framework relied on bulk microbial indicators to estimate cell culturability, the close agreement between bulk and *L. pneumophila*-specific culturable ratios in most samples supports the practical value of this proxy for quantitative interpretation. However, deviations observed in partial samples underscores the urgency to develop rapid methods to assess cell culturability in drinking water. Overcoming this biological measurement gap would further improve the speed, reliability, and operational value of routine *L. pneumophila* monitoring by molecular assay.

#### Notes

The authors declare no competing financial interest.

## Acknowledgments

This work was supported by the U.S. Environmental Protection Agency National Priorities Program (Grant No. 8406060). We also thank the participating water utility personnel for their assistance in this study.

## Reference

(1) Mercante Jeffrey W.; Winchell Jonas M. Current and Emerging Legionella Diagnostics for Laboratory and Outbreak Investigations. Clinical Microbiology Reviews 2015, 28 (1), 95–133. 10.1128/cmr.00029-14.

(2) Schwartz, T.; Hoffmann, S.; Obst, U. Formation of Natural Biofilms during Chlorine Dioxide and u.v. Disinfection in a Public Drinking Water Distribution System. Journal of Applied Microbiology 2003, *S5* (3), 591–601. 10.1046/j.1365-2672.2003.02019.x.

(3) Viasus, D.; Gaia, V.; Manzur-Barbur, C.; Carratalà, J. Legionnaires’ Disease: Update on Diagnosis and Treatment. Infectious Diseases and Therapy 2022, 11 (3), 973–986. 10.1007/s40121-022-00635-7.

(4) Lesnik, R.; Brettar, I.; Höfle, M. G. *Legionella* Species Diversity and Dynamics from Surface Reservoir to Tap Water: From Cold Adaptation to Thermophily. The ISME Journal 2016, 10 (5), 1064–1080. 10.1038/ismej.2015.199.

(5) Doleans Anne; Aurell Helena; Reyrolle Monique; Lina Gerard; Freney Jean; Vandenesch Francois; Etienne Jerome; Jarraud Sophie. Clinical and Environmental Distributions of Legionella Strains in France Are Different. Journal of Clinical Microbiology 2004, 42 (1), 458–460. 10.1128/jcm.42.1.458-460.2004.

(6) Yu, V. L.; Plouffe, J. F.; Pastoris, M. C.; Stout, J. E.; Schousboe, M.; Widmer, A.; Summersgill, J.; File, T.; Heath, C. M.; Paterson, D. L.; Chereshsky, A. Distribution of Legionella Species and Serogroups Isolated by Culture in Patients with Sporadic Community-Acquired Legionellosis: An International Collaborative Survey.

(7) Xi, H.; Ross, K. E.; Hinds, J.; Molino, P. J.; Whiley, H. Efficacy of Chlorine-Based Disinfectants to Control Legionella within Premise Plumbing Systems. Water Research 2024, *25S*, 121794. 10.1016/j.watres.2024.121794.

(8) Rasheduzzaman, M.; Singh, R.; Haas, C. N.; Gurian, P. L. Required Water Temperature in Hotel Plumbing to Control Legionella Growth. Water Research 2020, 182, 115943. 10.1016/j.watres.2020.115943.

(9) Clements, E.; Crank, K.; Nerenberg, R.; Atkinson, A.; Gerrity, D.; Hannoun, D. Quantitative Microbial Risk Assessment Framework Incorporating Water Ages with Legionella Pneumophila Growth Rates. Environ. Sci. Technol. 2024, 58 (15), 6540–6551. 10.1021/acs.est.4c01208.

(10) Lau, H. Y.; Ashbolt, N. J. The Role of Biofilms and Protozoa in Legionella Pathogenesis: Implications for Drinking Water. Journal of applied microbiology 2009, 107 (2), 368–378.

(11) Hamilton, K. A.; Prussin, A. J.; Ahmed, W.; Haas, C. N. Outbreaks of Legionnaires’ Disease and Pontiac Fever 2006–2017. Current environmental health reports 2018, 5 (2), 263–271.

(12) Garrison, L. E.; Kunz, J. M.; Cooley, L. A.; Moore, M. R.; Lucas, C.; Schrag, S.; Sarisky, J.; Whitney, C. G. Vital Signs: Deficiencies in Environmental Control Identified in Outbreaks of Legionnaires’ Disease—North America, 2000–2014. American Journal of Transplantation 2016, 1c (10), 3049–3058. 10.1111/ajt.14024.

(13) Kunz, J. M. Surveillance of Waterborne Disease Outbreaks Associated with Drinking Water—United States, 2015–2020. MMWR. Surveillance Summaries 2024, 73.

(14) Prussin II, A. J.; Schwake, D. O.; Marr, L. C. Ten Questions Concerning the Aerosolization and Transmission of Legionella in the Built Environment. Building and environment 2017, 123, 684–695.

(15) Hamilton, K. A.; Hamilton, M. T.; Johnson, W.; Jjemba, P.; Bukhari, Z.; LeChevallier, M.; Haas, C. N.; Gurian, P. L. Risk-Based Critical Concentrations of Legionella Pneumophila for Indoor Residential Water Uses. Environ. Sci. Technol. 2019, 53 (8), 4528–4541. 10.1021/acs.est.8b03000.

(16) Blatt, S. P.; Parkinson, M. D.; Pace, E.; Hoffman, P.; Dolan, D.; Lauderdale, P.; Zajac, R. A.; Melcher, G. P. Nosocomial Legionnaires’ Disease: Aspiration as a Primary Mode of Disease Acquisition. The American journal of medicine 1993, S5 (1), 16–22.

(17) Gleason, J. A.; Cohn, P. D. A Review of Legionnaires’ Disease and Public Water Systems–Scientific Considerations, Uncertainties and Recommendations. International Journal of Hygiene and Environmental Health 2022, 240, 113906.

(18) Code of Federal Regulations. National Primary Drinking Water Regulations; 1975. https://www.ecfr.gov/current/title-40/chapter-I/subchapter-D/part-141.

(19) U.S. Environmental Protection Agency. Surface Water Treatment Rules; 1989; Vol. 40 CFR Part 141, Subpart H. https://www.ecfr.gov/current/title-40/chapter-I/subchapter-D/part-141/subpart-H.

(20) 9260 INTRODUCTION TO DETECTING PATHOGENIC BACTERIA. In Standard Methods For the Examination of Water and Wastewater; Standard Methods for the Examination of Water and Wastewater; American Public Health Association, 2017. 10.2105/SMWW.2882.201.

(21) Centers for Disease Control and Prevention. Methods for Legionella Testing and Specimen Collection; Guidance; Centers for Disease Control and Prevention: Atlanta, GA.

(22) Walker, J. T.; McDermott, P. J. Confirming the Presence of Legionella Pneumophila in Your Water System: A Review of Current Legionella Testing Methods. Journal of AOAC International 2021, 104 (4), 1135–1147.

(23) Bartie, C.; Venter, S. N.; Nel, L. H. Identification Methods for Legionella from Environmental Samples. Water research 2003, 37 (6), 1362–1370.

(24) Lucas, C. E.; Taylor Jr, T. H.; Fields, B. S. Accuracy and Precision of Legionella Isolation by US Laboratories in the ELITE Program Pilot Study. Water research 2011, 45 (15), 4428–4436.

(25) Boczek, L. A.; Tang, M.; Formal, C.; Lytle, D.; Ryu, H. Comparison of Two Culture Methods for the Enumeration of Legionella Pneumophila from Potable Water Samples. Journal of Water and Health 2021, *1S* (3), 468–477. 10.2166/wh.2021.051.

(26) Spies, K.; Pleischl, S.; Lange, B.; Langer, B.; Hübner, I.; Jurzik, L.; Luden, K.; Exner, M. Comparison of the Legiolert^TM^/Quanti-Tray® MPN Test for the Enumeration of Legionella Pneumophila from Potable Water Samples with the German Regulatory Requirements Methods ISO 11731-2 and ISO 11731. International journal of hygiene and environmental health 2018, 221 (7), 1047–1053.

(27) Sartory, D. P.; Spies, K.; Lange, B.; Schneider, S.; Langer, B. Evaluation of a Most Probable Number Method for the Enumeration of Legionella Pneumophila from Potable and Related Water Samples. Letters in Applied Microbiology 2017, *c4* (4), 271–275. 10.1111/lam.12719.

(28) Rech, M. M.; Swalla, B. M.; Dobranic, J. K. Evaluation of Legiolert for Quantification of Legionella Pneumophila from Non-Potable Water. Current Microbiology 2018, 75 (10), 1282–1289. 10.1007/s00284-018-1522-0.

(29) Nazarian, E. J.; Bopp, D. J.; Saylors, A.; Limberger, R. J.; Musser, K. A. Design and Implementation of a Protocol for the Detection of Legionella in Clinical and Environmental Samples. Diagnostic microbiology and infectious disease 2008, *c2* (2), 125–132.

(30) Lee, J. V.; Lai, S.; Exner, M.; Lenz, J.; Gaia, V.; Casati, S.; Hartemann, P.; Lück, C.; Pangon, B.; Ricci, M. L.; Scaturro, M.; Fontana, S.; Sabria, M.; Sánchez, I.; Assaf, S.; Surman-Lee, S. An International Trial of Quantitative PCR for Monitoring Legionella in Artificial Water Systems. Journal of Applied Microbiology 2011, 110 (4), 1032–1044. 10.1111/j.1365-2672.2011.04957.x.

(31) Arrigo, I.; Serra, N.; Mariyam, L.; Sarvandani, M. M.; Tricoli, M. R.; Palermo, R.; Diquattro, O.; Gallina, G.; Di Carlo, P.; Firenze, A.; Palermo, M.; Giammanco, A.; Fasciana, T. M. A. Comparison between Legiolert and Real Time PCR in the Detection of Legionella Pneumophila from Environmental Water Samples. PLOS ONE 2025, 20 (12), e0336207. 10.1371/journal.pone.0336207.

(32) Sylvestre, É.; Rhoads, W. J.; Julian, T. R.; Hammes, F. Quantification of Legionella Pneumophila in Building Potable Water Systems: A Meta-Analysis Comparing qPCR and Culture-Based Detection Methods. PLOS Water 2025, 4 (1), e0000291. 10.1371/journal.pwat.0000291.

(33) Guillemet, T. A.; Lévesque, B.; Gauvin, D.; Brousseau, N.; Giroux, J.-P.; Cantin, P. Assessment of Real-time PCR for Quantification of Legionella Spp. in Spa Water. Letters in Applied Microbiology 2010, 51 (6), 639–644. 10.1111/j.1472-765X.2010.02947.x.

(34) Collins, S.; Stevenson, D.; Walker, J.; Bennett, A. Evaluation of Legionella Real-time PCR against Traditional Culture for Routine and Public Health Testing of Water Samples. Journal of Applied Microbiology 2017, 122 (6), 1692–1703. 10.1111/jam.13461.

(35) Joly Philippe; Falconnet Pierre-Alain; André Janine; Weill Nicole; Reyrolle Monique; Vandenesch François; Maurin Max; Etienne Jerome; Jarraud Sophie. Quantitative Real-Time Legionella PCR for Environmental Water Samples: Data Interpretation. Applied and Environmental Microbiology 2006, 72 (4), 2801–2808. 10.1128/AEM.72.4.2801-2808.2006.

(36) Scaturro Maria; Girolamo Antonietta; Bella Antonino; Rota Maria Cristina; Marella Anna Maria; Amari Rossana; Ingrassia Massimiliano; Barberis Daria; Romano Chiara; Borney Francesca; Damasco Florida; Buratto Maria Teresa; Fortin Monica; Caricato Gaetano; La Vecchia Giovanna; Casaburi Filomena; Dragone Melania; Cavallaro Mario; Tozzi Giovanna; Ceccarelli Elisabetta; Coniglio Maria Anna; Agodi Antonella; Coroneo Valentina; Sanna Adriana; Cristina Maria Luisa; Spagnolo Anna Maria; Cristino Sandra; Spiteri Simona; De Lellis Laura; De Mirto Daniela; Ferrari Fabio; Guiso Maria Giovanna; Foti Marina; Graziano Elisabetta; Franchi Marinella; Felice Antonella; Giammanco Anna; Fasciana Teresa; Grucci Annalisa; Ballarini Elena; Helfer Fabrizia; Corradi Ivan; Laganà Pasqualina; Facciolà Alessio; Mansi Antonella; Marcelloni Anna Maria; Marchesi Isabella; Bargellini Annalisi; Mari Marianna; Palmieri Sabina; Miana Paola; Della Sala Stefano; De Giglio Osvalda; Montagna Maria Teresa; Diella Giusy; Moschin Anna; Lugarini Giorgia; Ottaviano Claudio; Caprini Andrea; Rossi Anna Maria; Pagano Mariangela; Talini Mariella; Gestri Donella; Vespa Giovannella; Croce Carla; Viggiani Mariagabriella; Livi Silvia; Ricci Maria Luisa. Interlaboratory Comparison of Culture-and PCR-Based Methods for Legionella Pneumophila Detection in Drinking Water Samples. Applied and Environmental Microbiology 2025, *S1* (6), e00236-25. 10.1128/aem.00236-25.

(37) Monteiro, S. N.; Robalo, A. M.; Santos, R. J. Evaluation of Legiolert^TM^ for the Detection of Legionella Pneumophila and Comparison with Spread-Plate Culture and qPCR Methods. Current Microbiology 2021, 78 (5), 1792–1797. 10.1007/s00284-021-02436-6.

(38) Wellinghausen Nele; Frost Cathrin; Marre Reinhard. Detection of Legionellae in Hospital Water Samples by Quantitative Real-Time LightCycler PCR. Applied and Environmental Microbiology 2001, *c7* (9), 3985–3993. 10.1128/AEM.67.9.3985-3993.2001.

(39) He, H.; DiLoreto, S.; Yang, J.; Milne, P.; Impellitteri, C. A.; Stubbins, A.; Pieper, K.; Graham, K.; Huang, C.-H.; Pinto, A. Legionella and Mycobacterium Populations Exhibit Geographic Structuring across and within Drinking Water Systems. bioRxiv 2026, 2026.01.13.699378. 10.64898/2026.01.13.699378.

(40) Yang, J.; He, H.; DiLoreto, S.; Bian, K.; Phaneuf, J. R.; Milne, P.; Pieper, K.; Stubbins, A.; Huang, C.-H.; Graham, K. E.; Impellitteri, C. A.; Pinto, A. Interpretable Machine Learning Reveals Integrated Water Chemistry and Parameter-Specific Nonlinear Responses Shaping <em>Legionella</em> Spp. and <em>Mycobacterium</em> Spp. in Drinking Water. medRxiv 2026, 2026.04.23.26351579. 10.64898/2026.04.23.26351579.

(41) DiLoreto, S.; He, H.; Yang, J.; Milne, P.; Li, J.; Impellitteri, C. A.; Stubbins, A.; Pinto, A.; Huang, C.-H. Identification of Factors Influencing Variability in Disinfection Byproducts and Their Toxicity in Chlorinated and Chloraminated Drinking Water Distribution Systems across the United States. Environ. Sci. Technol. 2026, *c0* (1), 1241–1252. 10.1021/acs.est.5c12121.

(42) Nazarian, E. J.; Bopp, D. J.; Saylors, A.; Limberger, R. J.; Musser, K. A. Design and Implementation of a Protocol for the Detection of Legionella in Clinical and Environmental Samples. Diagnostic Microbiology and Infectious Disease 2008, *c2* (2), 125–132. 10.1016/j.diagmicrobio.2008.05.004.

(43) Schmidt, M. P.; Martínez, C. E. Supramolecular Association Impacts Biomolecule Adsorption onto Goethite. Environ. Sci. Technol. 2018, 52 (7), 4079–4089. 10.1021/acs.est.7b06173.

(44) Sudarshan, A. S.; Dai, Z.; Gabrielli, M.; Oosthuizen-Vosloo, S.; Konstantinidis, K. T.; Pinto, A. J. New Drinking Water Genome Catalog Identifies a Globally Distributed Bacterial Genus Adapted to Disinfected Drinking Water Systems. Environ. Sci. Technol. 2024, 58 (37), 16475–16487. 10.1021/acs.est.4c05086.

(45) Robert, C. P.; Casella, G.; Casella, G. Monte Carlo Statistical Methods; Springer, 2004; Vol. 2.

(46) Taper, M. L.; Lele, S. R.; Ponciano, J. M.; Dennis, B.; Jerde, C. L. Assessing the Global and Local Uncertainty of Scientific Evidence in the Presence of Model Misspecification. Frontiers in Ecology and Evolution 2021, *Volume S-*2021. 10.3389/fevo.2021.679155.

(47) Urquizo, J.; Calderón, C.; James, P. Using a Local Framework Combining Principal Component Regression and Monte Carlo Simulation for Uncertainty and Sensitivity Analysis of a Domestic Energy Model in Sub-City Areas. Energies 2017, 10 (12), 1986. 10.3390/en10121986.

(48) Goovaerts, P. Geostatistical Analysis of Disease Data: Estimation of Cancer Mortality Risk from Empirical Frequencies Using Poisson Kriging. International Journal of Health Geographics 2005, 4 (1), 31. 10.1186/1476-072X-4-31.

(49) Ganser, G. H.; Hewett, P. An Accurate Substitution Method for Analyzing Censored Data. Journal of Occupational and Environmental Hygiene 2010, 7 (4), 233–244. 10.1080/15459621003609713.

(50) Ma, J.-X.; Wang, X.; Pan, Y.-R.; Wang, Z.-Y.; Guo, X.; Liu, J.; Ren, N.-Q.; Butler, D. Data-Driven Systematic Analysis of Waterborne Viruses and Health Risks during the Wastewater Reclamation Process. Environmental Science and Ecotechnology 2024, *1S*, 100328.

(51) LeChevallier, M. W. Occurrence of Culturable Legionella Pneumophila in Drinking Water Distribution Systems. AWWA Water Science 2019, 1 (3), e1139. 10.1002/aws2.1139.

(52) Putri, R. E.; Kim, L. H.; Farhat, N.; Felemban, M.; Saikaly, P. E.; Vrouwenvelder, J. S. Evaluation of DNA Extraction Yield from a Chlorinated Drinking Water Distribution System. PLOS ONE 2021, *1c* (6), e0253799. 10.1371/journal.pone.0253799.

(53) Putri, R. E.; Vrouwenvelder, J. S.; Farhat, N. Enhancing the DNA Yield Intended for Microbial Sequencing from a Low-Biomass Chlorinated Drinking Water. Frontiers in Microbiology 2024, *Volume 15-*2024.

(54) Delgado-Viscogliosi Pilar; Solignac Lydie; Delattre Jean-Marie. Viability PCR, a Culture-Independent Method for Rapid and Selective Quantification of Viable Legionella Pneumophila Cells in Environmental Water Samples. Applied and Environmental Microbiology 2009, 75 (11), 3502–3512. 10.1128/AEM.02878-08.

(55) Guo, L.; Xiao, X.; Chabi, K.; Zhang, Y.; Li, J.; Yao, S.; Yu, X. Occurrence of Viable but Non-Culturable (VBNC) Pathogenic Bacteria in Tap Water of Public Places. Frontiers of Environmental Science & Engineering 2023, 18 (3), 35. 10.1007/s11783-024-1795-4.

(56) Alleron, L.; Merlet, N.; Lacombe, C.; Frère, J. Long-Term Survival of Legionella Pneumophila in the Viable But Nonculturable State After Monochloramine Treatment. Current Microbiology 2008, 57 (5), 497–502. 10.1007/s00284-008-9275-9.

(57) Hu, Z.; Bai, X. Self-Repair and Resuscitation of Viable Injured Bacteria in Chlorinated Drinking Water: Achromobacter as an Example. Water Research 2023, 245, 120585. 10.1016/j.watres.2023.120585.

(58) Ramseier, M. K.; von Gunten, U.; Freihofer, P.; Hammes, F. Kinetics of Membrane Damage to High (HNA) and Low (LNA) Nucleic Acid Bacterial Clusters in Drinking Water by Ozone, Chlorine, Chlorine Dioxide, Monochloramine, Ferrate(VI), and Permanganate. Water Research 2011, 45 (3), 1490–1500. 10.1016/j.watres.2010.11.016.

(59) Bairoliya, S.; Goel, A.; Mukherjee, M.; Koh Zhi Xiang, J.; Cao, B. Monochloramine Induces Release of DNA and RNA from Bacterial Cells: Quantification, Sequencing Analyses, and Implications. Environ. Sci. Technol. 2022, *5c* (22), 15791–15804. 10.1021/acs.est.2c06632.

(60) Sakcham, B.; Kumar, A.; Cao, B. Extracellular DNA in Monochloraminated Drinking Water and Its Influence on DNA-Based Profiling of a Microbial Community. Environ. Sci. Technol. Lett. 2019, c (5), 306–312. 10.1021/acs.estlett.9b00185.

(61) Donohue, I.; Irvine, K. Quantifying Variability within Water Samples: The Need for Adequate Subsampling. Water research 2008, 42 (1–2), 476–482.

(62) LeChevallier, M. W. Monitoring Distribution Systems for Legionella Pneumophila Using Legiolert. AWWA Water Science 2019, 1 (1), e1122. 10.1002/aws2.1122.

(63) Hirsh, M.; Baron, J. L.; Mietzner, S.; Rihs, J. D.; Stout, J. E. Cross-reactivity of the IDEXX Legiolert Method with Other Gram-negative Bacteria and Waterborne Pathogens Leads to False-positive Assay Results. Letters in applied microbiology 2021, 72 (6), 750–756.

(64) Van Thienen, P.; Vreeburg, J. H. G. Turbulent Processes in Drinking Water Distribution. In Water Distribution Systems Analysis 2010; American Society of Civil Engineers: Tucson, Arizona, United States, 2011; pp 635–642. 10.1061/41203(425)60.

(65) Peng Zhangjie; Stovin Virginia; Guymer Ian. Radial Mixing in Steady and Accelerating Pipe Flows. Journal of Hydraulic Engineering 2024, 150 (6), 04024047. 10.1061/JHEND8.HYENG-14071.

(66) Van Thienen, P.; Vreeburg, J. H. G.; Blokker, E. J. M. Radial Transport Processes as a Precursor to Particle Deposition in Drinking Water Distribution Systems. Water research 2011, 45 (4), 1807–1817.

(67) Bižić-Ionescu, M.; Ionescu, D.; Grossart, H.-P. Organic Particles: Heterogeneous Hubs for Microbial Interactions in Aquatic Ecosystems. Frontiers in Microbiology 2018, *S*, 2569.

(68) Droppo, I. G.; Liss, S. N.; Williams, D.; Nelson, T.; Jaskot, C.; Trapp, B. Dynamic Existence of Waterborne Pathogens within River Sediment Compartments. Implications for Water Quality Regulatory Affairs. Environ. Sci. Technol. 2009, 43 (6), 1737–1743. 10.1021/es802321w.

(69) Bédard, E.; Laferrière, C.; Déziel, E.; Prévost, M. Impact of Stagnation and Sampling Volume on Water Microbial Quality Monitoring in Large Buildings. PLOS ONE 2018, 13 (6), e0199429. 10.1371/journal.pone.0199429.

(70) Prévost, M.; Rompré, A.; Baribeau, H.; Coallier, J.; Lafrance, P. Service Lines: Their Effect on Microbiological Quality. Journal AWWA 1997, 8S (7), 78–92. 10.1002/j.1551-8833.1997.tb08261.x.

(71) Buchberger, S. G.; Lee, Y.; Bloom, G.; Rolf, B. Dispersion of Mass in Intermittent Laminar Flow through Pipe. Water industry systems: modeling and optimization applications 1999, 1, 89–101.

(72) Buchberger, S. G.; Lee, Y. Evidence Supporting the Poisson Pulse Hypothesis for Residential Water Demands. Water industry systems: Modelling and optimization applications 1999, 215–227.

(73) Tzatchkov Velitchko G.; Aldama Alvaro A.; Arreguin Felipe I. Advection-Dispersion-Reaction Modeling in Water Distribution Networks. Journal of Water Resources Planning and Management 2002, 128 (5), 334–342. 10.1061/(ASCE)0733-9496(2002)128:5(334).

(74) Zhang, C.; Lu, J. Legionella: A Promising Supplementary Indicator of Microbial Drinking Water Quality in Municipal Engineered Water Systems. Frontiers in environmental science 2021, *S*, 684319.

(75) Duda, S.; Baron, J. L.; Wagener, M. M.; Vidic, R. D.; Stout, J. E. Lack of Correlation between Legionella Colonization and Microbial Population Quantification Using Heterotrophic Plate Count and Adenosine Triphosphate Bioluminescence Measurement. Environmental Monitoring and Assessment 2015, 187 (7), 393. 10.1007/s10661-015-4612-5.

(76) Shen, Y.; Huang, C.; Lin, J.; Wu, W.; Ashbolt, N. J.; Liu, W.-T.; Nguyen, T. H. Effect of Disinfectant Exposure on Legionella Pneumophila Associated with Simulated Drinking Water Biofilms: Release, Inactivation, and Infectivity. Environ. Sci. Technol. 2017, 51 (4), 2087–2095. 10.1021/acs.est.6b04754.

(77) Cambronne, E. D.; Ayres, C.; Dowdell, K. S.; Lawler, D. F.; Saleh, N. B.; Kirisits, M. J. Protozoan-Priming and Magnesium Conditioning Enhance Legionella Pneumophila Dissemination and Monochloramine Resistance. Environ. Sci. Technol. 2023, 57 (40), 14871–14880. 10.1021/acs.est.3c04013.

(78) Turetgen, I. Induction of Viable but Nonculturable (VBNC) State and the Effect of Multiple Subculturing on the Survival ofLegionella Pneumophila Strains in the Presence of Monochloramine. Annals of Microbiology 2008, 58 (1), 153–156. 10.1007/BF03179460.

(79) Alleron, L.; Khemiri, A.; Koubar, M.; Lacombe, C.; Coquet, L.; Cosette, P.; Jouenne, T.; Frere, J. VBNC Legionella Pneumophila Cells Are Still Able to Produce Virulence Proteins. Water research 2013, 47 (17), 6606–6617.

