## Supplemental information for "Accounting for DNA Recovery and Cell Culturability Enhances Quantitative Compatibility of Molecular and Legiolert Assays for *Legionella pneumophila*"

*Corresponding Author*

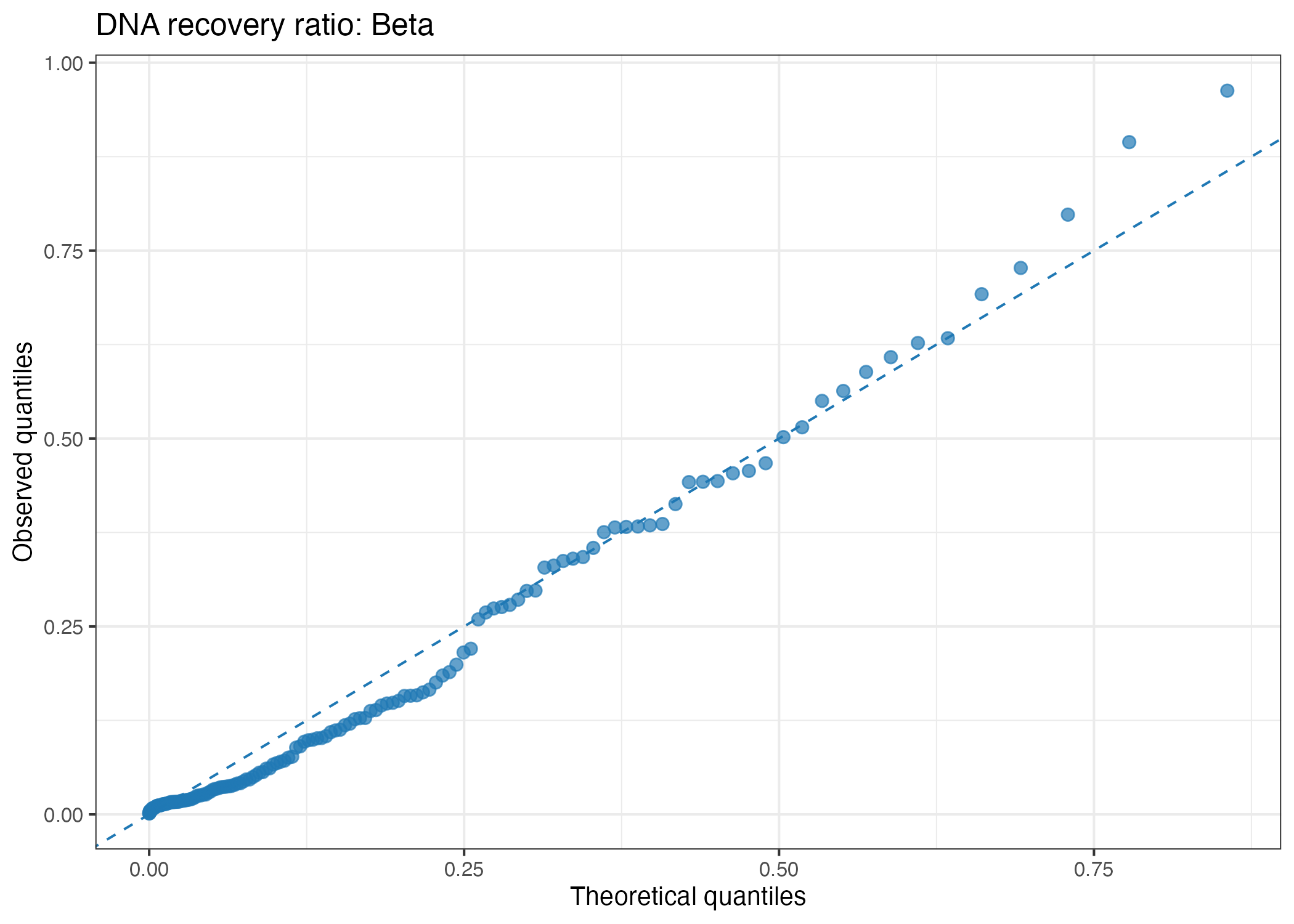

**Figure S1.** Q–Q plot of the beta-distribution fit for the DNA recovery ratio using the median MAG-derived genome size of 3.07 Mbp.

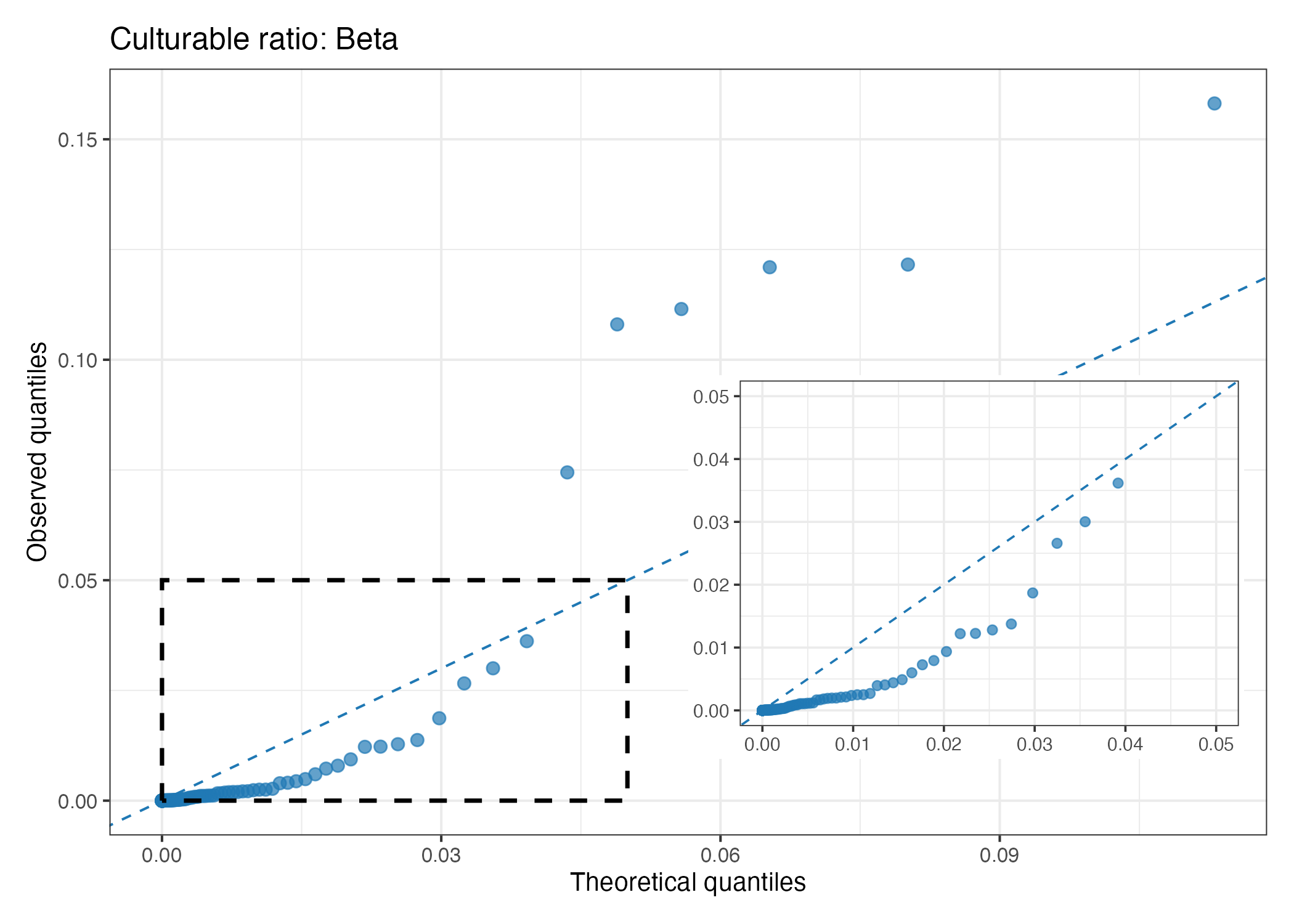

**Figure S2.** Q–Q plot of the beta-distribution fit for the bulk culturable ratio using the median MAG-derived genome size of 3.07 Mbp.

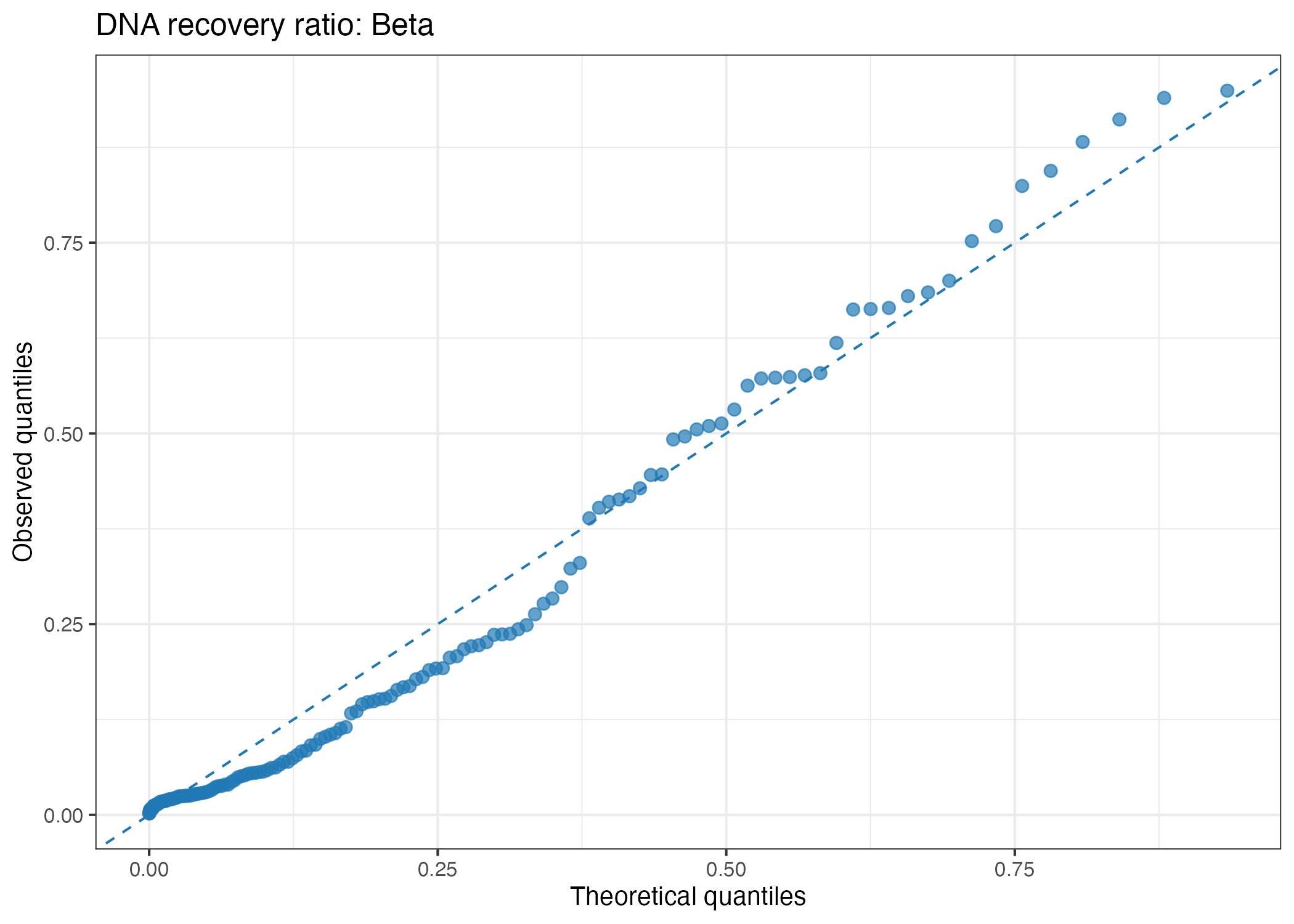

**Figure S3.** Q–Q plot of the beta-distribution fit for the DNA recovery ratio using the Q1 MAG-derived genome size 2.05 Mbp.

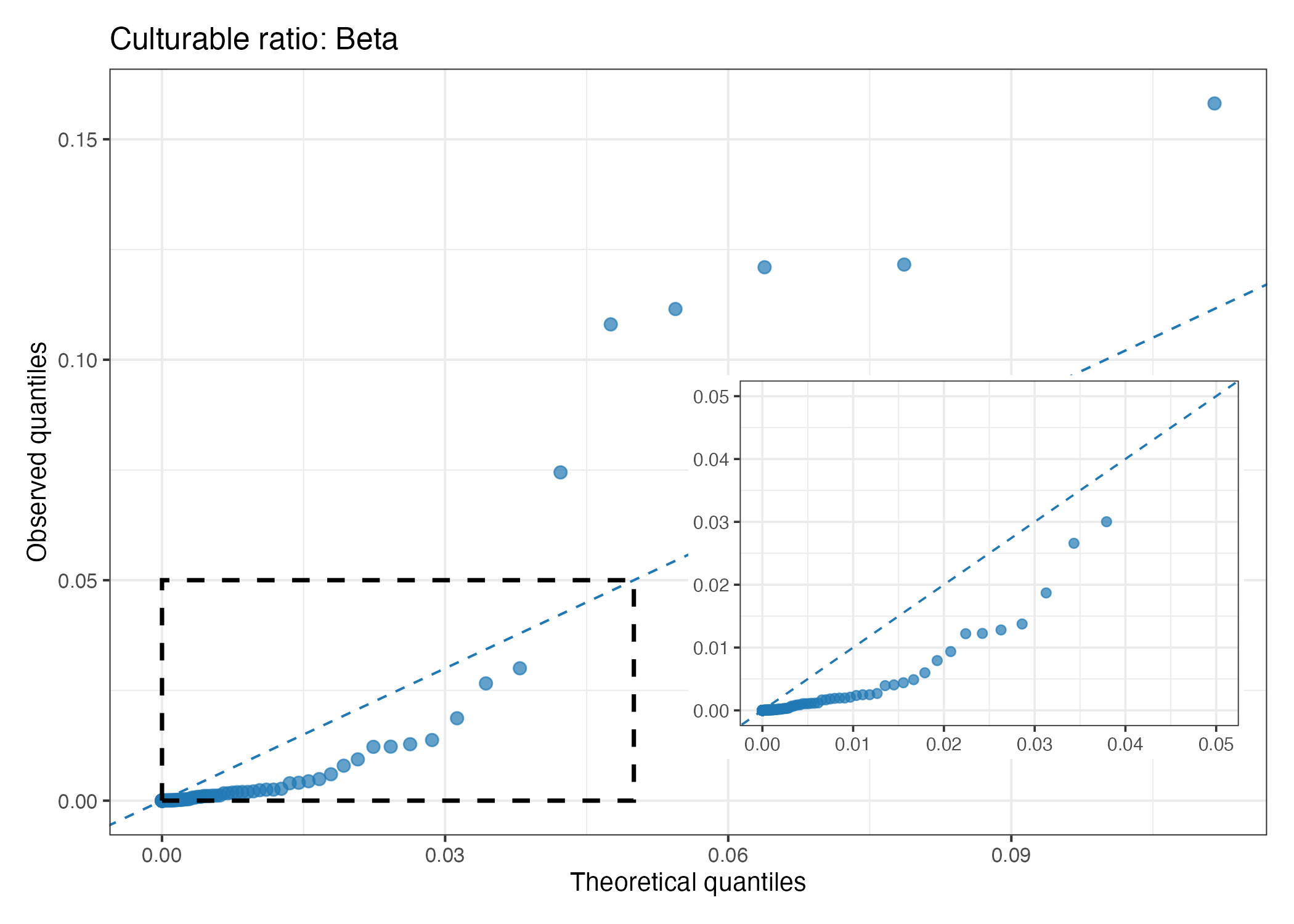

**Figure S4.** Q–Q plot of the beta-distribution fit for the bulk culturable ratio using the Q1 MAG-derived genome size 2.05 Mbp.

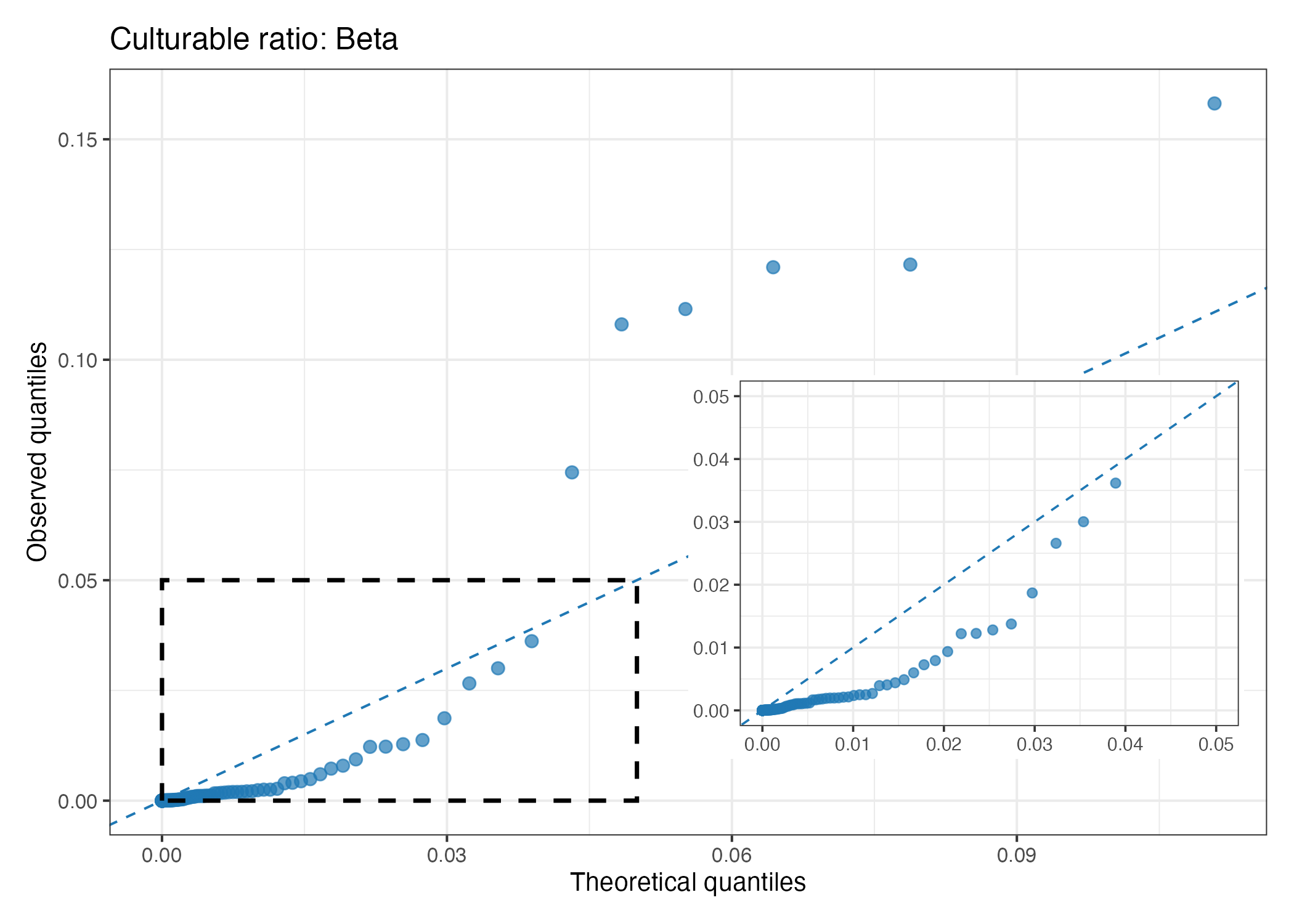

**Figure S5.** Q–Q plot of the beta-distribution fit for the bulk culturable ratio using the Q3 MAG-derived genome size 4.25 Mbp.

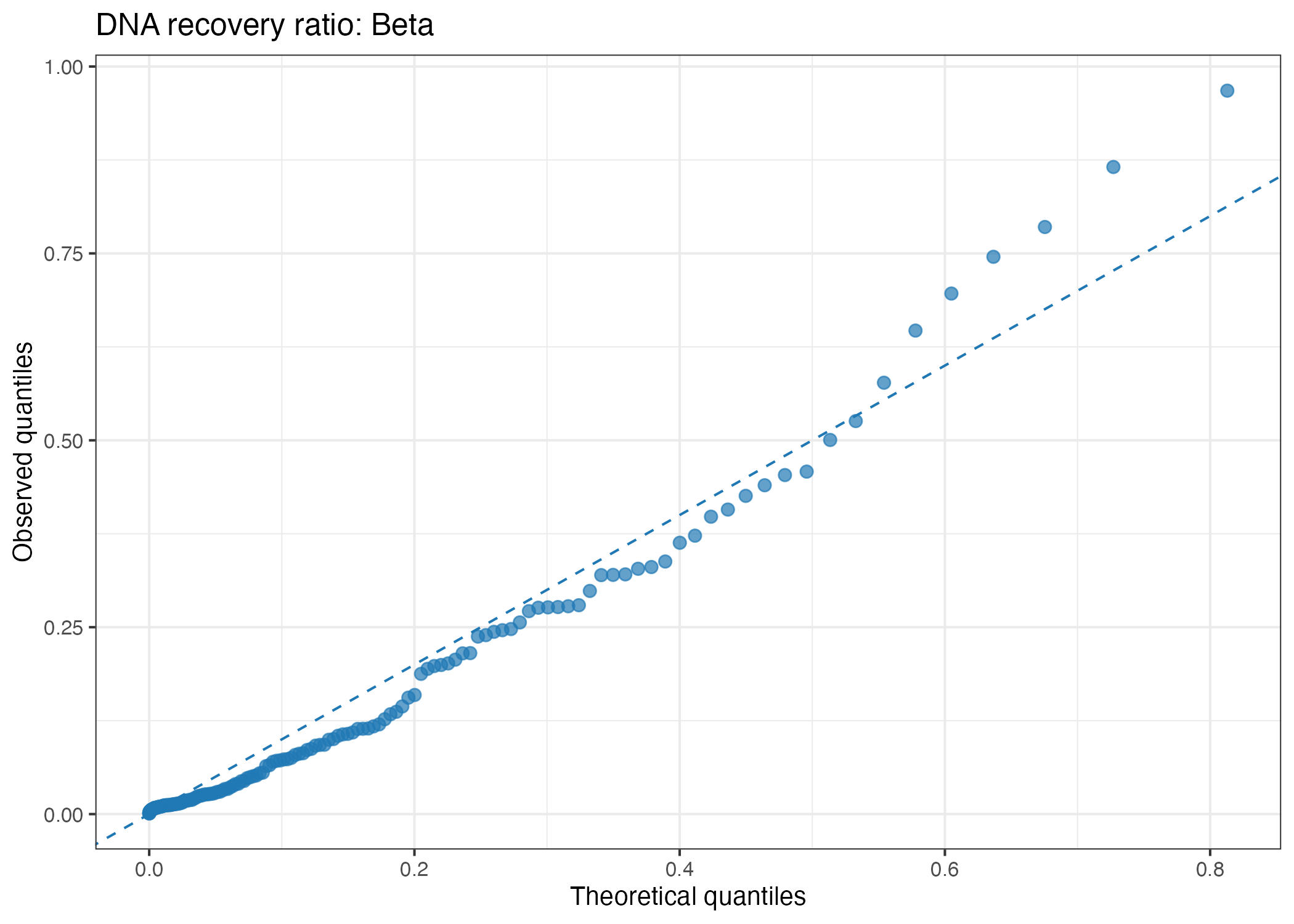

**Figure S6.** Q–Q plot of the beta-distribution fit for the DNA recovery ratio using the Q3 MAG-derived genome size 4.25 Mbp.

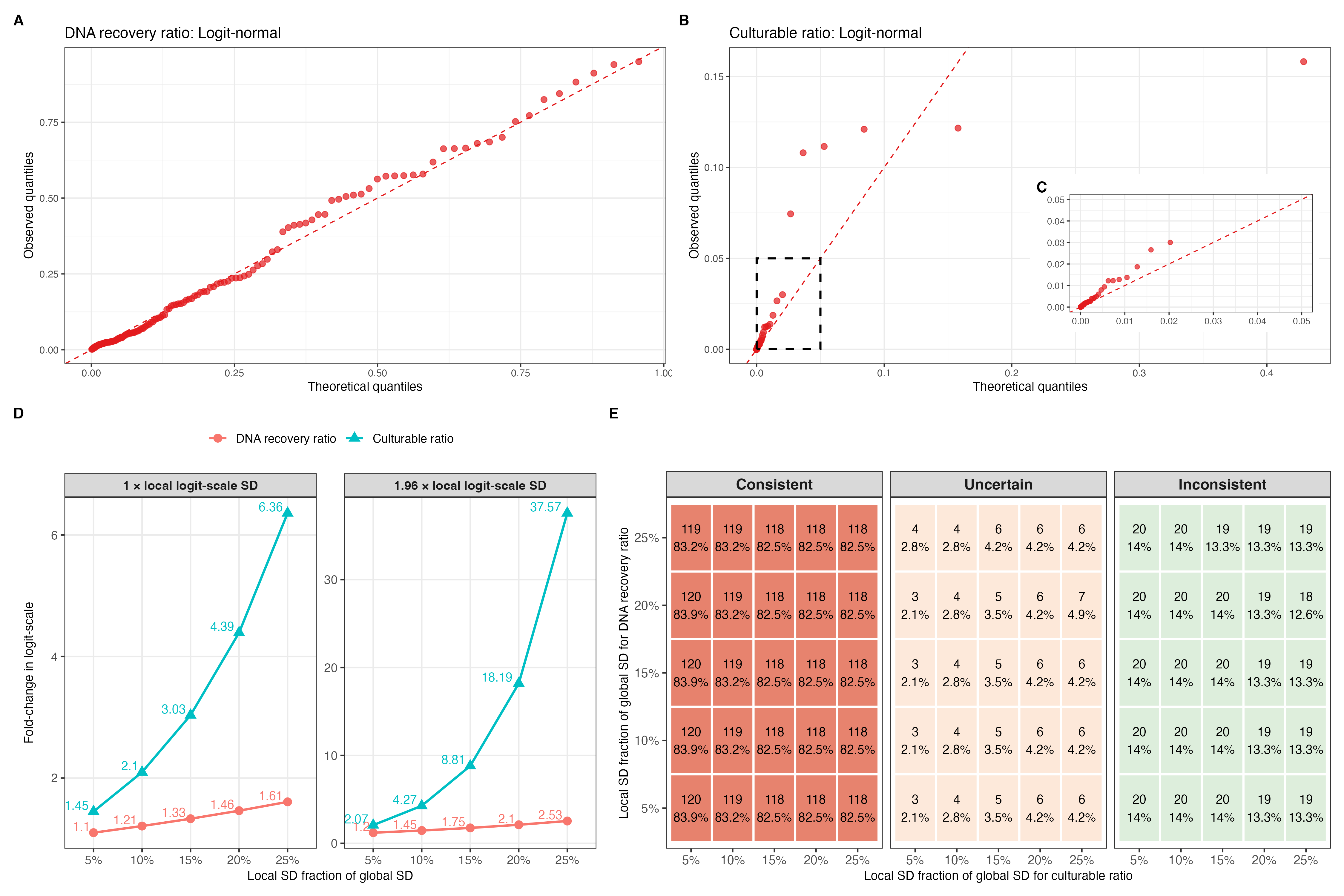

**Figure S7.** Monte Carlo parameterization and sensitivity analysis using the Q1 MAG-derived genome size of 2.05 Mbp. Q–Q plots of the logit-normal fits for the (A) DNA recovery ratio and (B) bulk culturable ratio; the inset in (C) shows the low-ratio region for the bulk culturable ratio. (D) Variation in logit-scale corresponding to 1 × and 1.96 × local logit-scale standard deviation for the DNA recovery ratio and bulk culturable ratio across local standard deviation fractions. (E) Consistency classifications across combinations of DNA-recovery and bulk-culturable-ratio local standard deviation fractions.

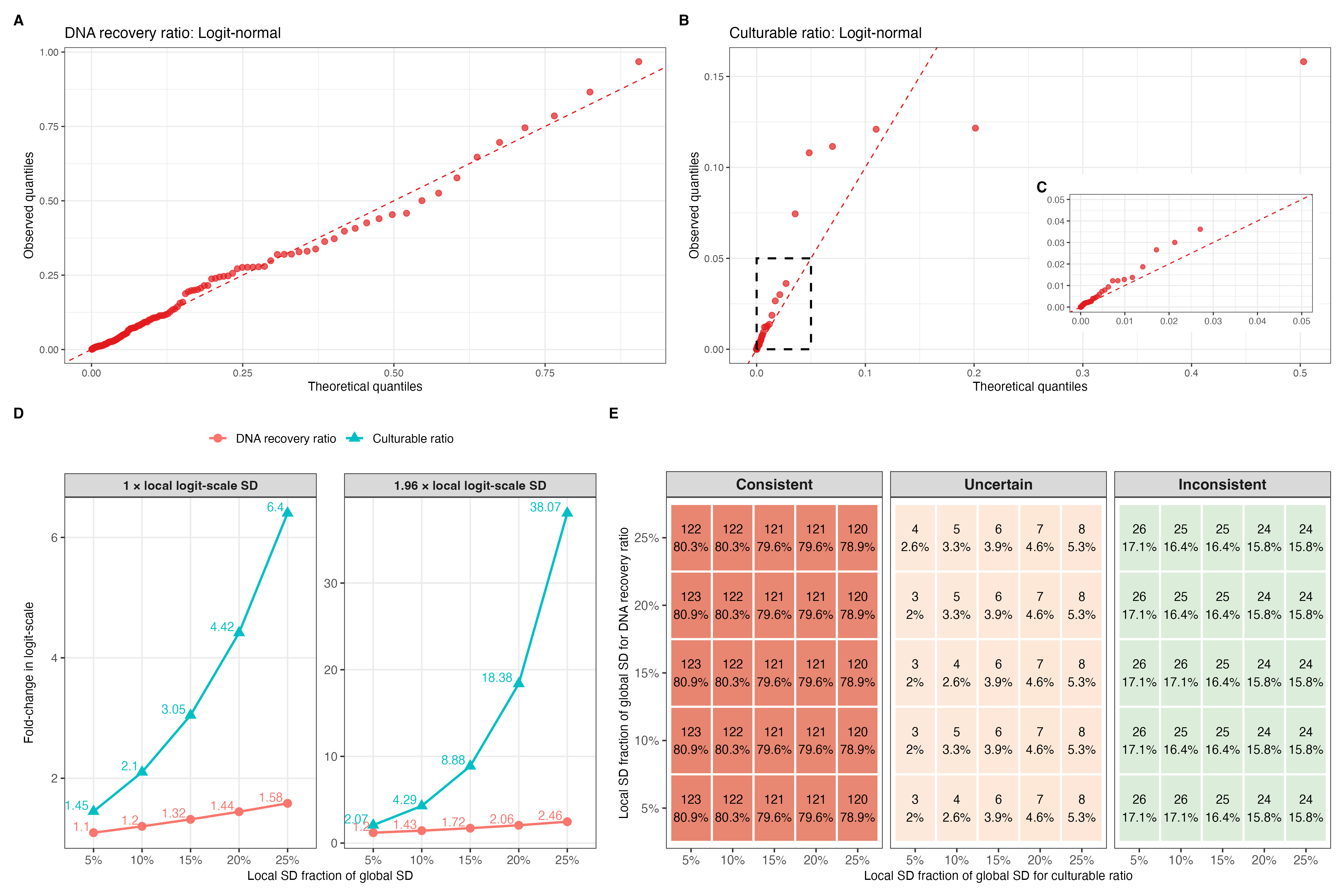

**Figure S8.** Monte Carlo parameterization and sensitivity analysis using the Q3 MAG-derived genome size of 4.25 Mbp. Q–Q plots of the logit-normal fits for the (A) DNA recovery ratio and (B) bulk culturable ratio; the inset in (C) shows the low-ratio region for the bulk culturable ratio. (D) Variation in logit-scale corresponding to 1 × and 1.96 × local logit-scale standard deviation for the DNA recovery ratio and bulk culturable ratio across local standard deviation fractions. (E) Consistency classifications across combinations of DNA-recovery and bulk-culturable-ratio local standard deviation fractions.

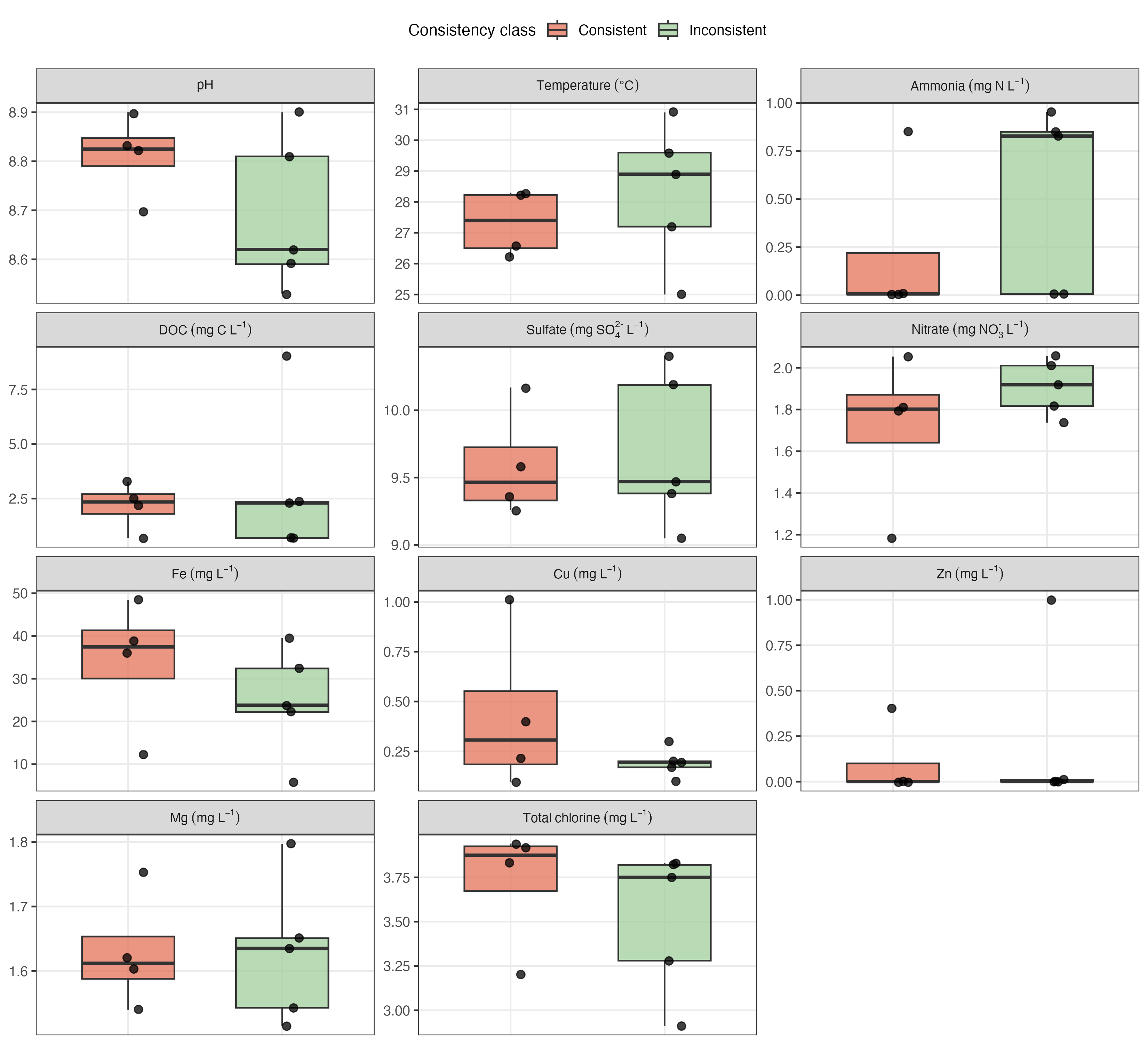

**Figure S9.** Physicochemical parameters of consistent and inconsistent Utility 5 finished drinking water samples from Rounds 2 and 3.
